# Internal evolutionary conflicts: a conceptual synthesis and mathematical primer

**DOI:** 10.64898/2026.07.16.739017

**Authors:** Gaurav S. Athreya, Ananda Shikhara Bhat, J. Arvid Ågren, E. Yagmur Erten, Thomas A. Keaney

## Abstract

Internal evolutionary conflicts arise when elements within an organism have diverging fitness interests. Examples range from meiotic drive and cytoplasmic male sterility to transposable elements and supernumerary B chromosomes. While once treated as genetic curiosities, they are now seen as widespread and major drivers of eukaryotic genome evolution. Yet their study remains fragmented, with no clear entry point not only for those who wish to gain an overview of theoretical advances, or those who wish to construct models of their own. Here, we discuss ways in which internal evolutionary conflicts have been modelled and develop a common population genetic framework for building such models. The framework provides explicit criteria for what counts as conflict, distinguishing it from fitness trade-offs, and formalises how and when internal conflicts arise. By treating different cases within the same structure, it shows that these diverse phenomena share a common logic.

## Introduction

Not all parts of an organism act in its best interest. Insect endosymbionts can kill or sterilise their male hosts, some genes benefit from helping kin to a far greater extent than others, while others still kill all of an individual’s gametes that they themselves aren’t found within. Despite this being clear for a long time (Gini, 1908; Gershenson, 1928; Lewis, 1941; Östergren, 1945), for much of the history of biology, internal evolutionary conflicts have been treated more as items in a Darwinian cabinet of curiosities than as central to evolutionary theory (Burt and Trivers (2006), pp.12-16; Rice (2013), p. 218; Ågren and Clark (2018), p.3). This picture has been shifting thanks to a close partnership between experiment and theory, with punctuated jumps fueled by a series of influential synthetic reviews. The result is an appreciation of internal evolutionary conflict as a rigorous branch of evolutionary biology, with some going so far as to describe it as essential to understanding all aspects of genetics (Rice, 2013).

The growth of the field can be traced back to an observation that some genes violate Mendel’s law of fair segregation (Sandler and Novitski, 1957). This discovery, coined meiotic drive, motivated the development of population genetic models, built to understand how distortions away from Mendelian transmission affect the evolutionary process (e.g. Bruck, 1957; Lewontin and Dunn, 1960; Charlesworth and Charlesworth, 1983). The breadth of internal evolutionary conflict was expanded by Cosmides and Tooby (1981), who applied a gene’s-eye view of evolution (Dawkins, 1976; Ågren, 2021) to derive a central principle: internal evolutionary conflicts often arise because not all genes are governed by the same inheritance rules. Behavioural ecologists, coming from a tradition of modelling animals as fitness-maximising agents, extended their inclusive fitness and game theoretic approaches to genes, adding mathematical precision to Cosmides and Tooby (1981)’s central logic. In addition to conflict produced by distorted segregation of alleles, they identified that within-organism conflicts could also manifest over sex allocation, as the sexes are not equally ‘valuable’ for allele propagation across the entirety of the genome (Hamilton, 1967). They further identified within-organism conflict over how individuals act towards genealogical relatives, as the degree of relatedness and the value a given relative has for allele propagation is also not consistent across the genome (Hamilton, 1972; Haig and Westoby, 1989).

The term ‘selfish genetic elements’ appeared in the late 1980s (Werren et al., 1988a), which was also the first attempt to discuss different kinds of genetic conflicts together as one common phenomenon. This was apt, as the catalogue of selfish elements had swollen, with often serendipitous discoveries illustrating that there are a diverse array of mechanisms that allow a gene to be ‘selfish’ (Hurst et al., 1996). By 2006, the field had its first book-length treatise, which exemplified the study of internal evolutionary conflicts as one of close dialogue between theory and experiment (Burt and Trivers, 2006). The shift from curiosity to mainstay was completed with an appreciation that internal evolutionary conflicts could be harnessed as biological control agents: the development of synthetic drive systems, designed to spread harmful alleles through populations at the expense of organismal level function, added an applied layer of interest to the literature (Champer et al., 2016; Burt and Koufopanou, 2004; Esvelt and Gemmell, 2017).

Yet, for all this progress, the breadth of empirical examples and theoretical approaches in the study of internal evolutionary conflicts can now make the field seem unwieldy, especially for those seeking to develop new theory. While there have been efforts to develop a unified mathematical framework (Gardner and Úbeda, 2017; Scott and West, 2019), it is not pedagogically clear how one can apply these frameworks to produce their own theoretical predictions. In particular, there is no gentle, self-contained introduction for an aspiring theoretician (or enthusiastic empiricist) that is general enough to guide one through the first steps of modelling their conflict of interest. Here, in the spirit of the close dialogue between theoreticians and empiricists that has shaped the field, we offer such an introductory primer for using a population genetic framework to build and analyse models of internal evolutionary conflict. In the next section, we begin by constructing a definition of internal evolutionary conflicts.

While certain phenomena are natural fits, others, although often invoked as internal evolutionary conflicts, are not well described by our approach. We then present an example mathematical model for each of the three forms of conflict that fall within our definition of internal evolutionary conflict. We choose models of tent-pole biological examples, well known to researchers of conflict, and show that learning the population genetic approach equips new theoreticians with the skills to tackle all types of internal evolutionary conflict. Throughout, we complement our main models with supplementary extensions, designed to deliver a deeper dive should the reader get interested. Lastly, we review ways in which these simple models have been extended for different biologically relevant purposes, and end by contrasting this framework with other existing approaches.

### What is an internal evolutionary conflict?

The study of internal evolutionary conflicts is complicated by terminology with multiple meanings (Gardner and Úbeda, 2017; Clarke, 2025). Our aim is not to resolve these debates over terminology. Rather, we begin by first setting out the criteria that serve as rules for deciding whether a given biological phenomenon counts as an internal evolutionary conflict in our framework.

First, our interaction of interest must be **internal** to one of the agents involved. All lower-level agents must be obligately contained within higher-level agents, in such a way that selective pressures affecting the ‘lower-level’ agent need not correlate with pressures that affect the encapsulating, ‘higher-level’ agent. For example, if a genome consists of elements that replicate normally, and in addition, elements that replicate faster and at the cost of organismal fitness, the faster replicators are favoured within each organism, but organisms without many fast replicators eventually become more common. This process can be described by the notion of multi-level selection (Okasha, 2006). Predator–prey interactions, for instance, are evolutionary conflicts (*sensu* Queller and Strassmann (2018)), but not internal since the prey is not always obligately contained inside the predator or vice versa. Note that interactions can be “internal” (i.e. involving multi-level selection) without necessarily requiring that the lower-level agent be exclusively vertically transmitted between higher-level agents. In particular, horizontal or oblique transmission of lower-level agents is permissible, but we *do* require that a lower-level agent cannot persist independently when not contained within the higher-level agent.

Second, the conflict must be **evolutionary** for all parties involved (*i.e*. there should be a co-evolutionary process). In other words, the consequences of the interactions between two parties are inherited **across generations by both parties**. As a consequence, conflicts such as those between an individual and a cancer it harbours are not evolutionary conflicts in our framework, since the consequences of the interaction are not inherited across generations on one side – that of the cancer. We provide a more detailed explanation of our rationale for excluding cancer from our framework in Box 1. If a eusocial colony is viewed as a superorganism, the same logic dictates that conflicts between a sterile worker and the colony cannot be viewed as an evolutionary conflict in our framework, since evolutionary consequences are not inherited across generations on the worker side. In particular, though non-heritable mutations that affect worker phenotypes can cause adaptive changes in colony composition and structure (Molet et al., 2012), this is analogous to the adaptive consequences of (non-inherited) somatic mutations in multicellular organisms (Majic et al., 2022), since different worker phenotypes are different cell or tissue types in the superorganism analogy.

Third, the interaction must involve a genuine **conflict** between multiple agents. By conflict we mean (at least) two distinct parties that influence a shared trait, phenotype, or outcome but have opposing interests in how it is expressed (Queller and Strassmann, 2018; Ågren et al., 2024). For instance, a game of chess involves a conflict because both players have (partial) control over the state of the board but have opposing interests. By this definition, a homing endonuclease that increases its own frequency without affecting organismal fitness (Burt and Koufopanou, 2004) is not a conflict but a commensal interaction. Similarly, though the components of the immune system within an organism can be thought of as being under the influence of evolutionary forces, there is no conflict at play because the fitness interests of, say, a white blood cell, are identical to that of the organism. Auto-immune disorders are thus not a case of evolutionary conflict but instead a case of loss of function/homeostasis.

A subtle point deserves mention here. Phenomena like antagonistic pleiotropy (Williams, 1957; Hedrick, 1999) and intra-locus sexual conflict (Haldane, 1962; Bonduriansky and Chenoweth, 2009) are not conflicts in our framework but rather trade-offs, since a gene’s expression in one context (young/female) changes at the expense of its expression in another context (old/male) — the two contexts are analogous to different environments, and a single party, the focal gene, simply has different fitness optima in the two environments (Cutter, 2023). While these trade-offs thus do not qualify as conflicts, there may indeed be internal evolutionary conflict over how a trade-off is resolved: alleles that are differently inherited (e.g. mitochondrial vs. nuclear) may have a different evolutionary interest in how this trade-off is balanced – even if they have identical fitness effects across two classes (e.g. male and female). In the next Section, we will show that an X-linked sexually antagonistic allele, i.e. one burdened by a fitness trade-off between the sexes, can invade even if it causes fitness costs so high in males that an autosomal version of the same allele (with identical fitness effects in males and females) would be pushed to extinction (Frank and Crespi, 2011).

**Box 1: Is cancer an internal evolutionary conflict?**

Cancer is a disease that arose with multicellularity (Aktipis et al., 2015), and the idea that it could be viewed as a “cell rebellion against the central authority” is more than a century old (Snow (1893), p. 4). At first sight, cancer may therefore appear to be a quintessential example of an internal evolutionary conflict: a conflict between the multicellular host and its non-cooperative cells (Laplane et al., 2025). On closer inspection, however, the picture is more complicated. In phenomena such as meiotic drive or sexratio distortion, the conflicting agents are inherited across multiple organismal generations and therefore co-evolve, fulfilling all three of our criteria for an internal evolutionary conflict (Fig 1). Tumour cells, by contrast, face an evolutionary dead end (Gardner, 2015; Okasha, 2024). The death of the host also results in the death of the tumour, so tumour cells do not meet the requirement that a conflict be evolutionary (see also pp. 41-43 of Shpak and Lu (2016)). They cannot co-evolve with the host, nor can they increase their own transmission across host generations.

**Figure 1.**
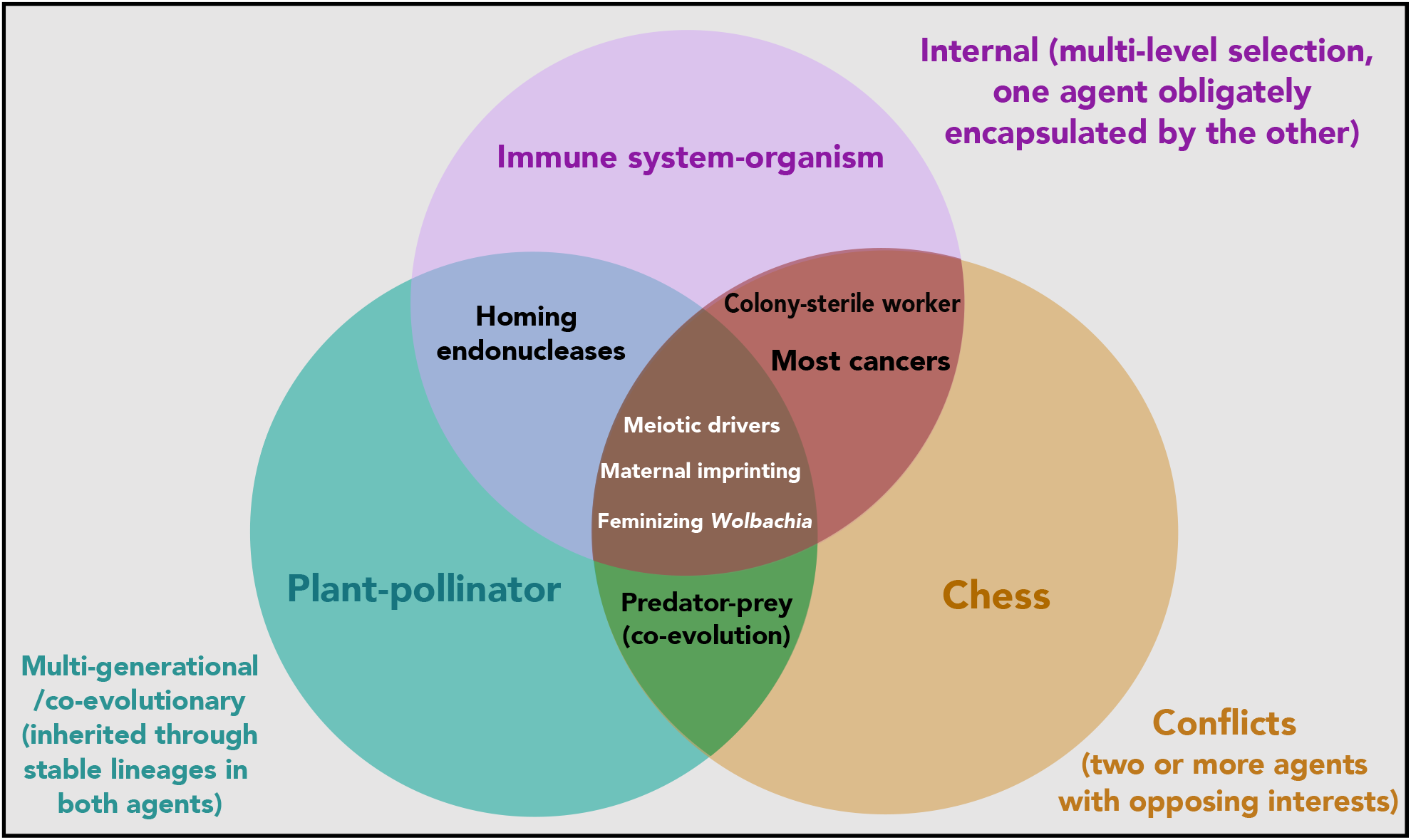
A classification scheme for inter-agent interactions illustrating what we mean by an internal evolutionary conflict. **Alt text**: Venn diagram involving 3 sets labelled “Internal”, “Multi-generational/Evolutionary”, and “Conflicts”. Each subset has an example, and almost all examples are mentioned in the main text.

Cancer, as a phenomenon, nonetheless has evolutionary consequences. For example, organisms that live with cancer risk evolve defences against it (Compton et al., 2025). In this case, however, what exerts a selective pressure is not a particular cancer but the phenomenon of cancer itself. In this respect, cancer is more akin to deleterious somatic mutations, both arise repeatedly in different hosts and in different ways, and selection acts to reduce their occurrence. By contrast, when a suppressor allele evolves in response to meiotic drive, it combats a specific drive element that is itself transmitted across generations, rather than the general phenomenon of drive.

Transmissible cancers are a clear exception to the arguments above. These tumour lineages not only proliferate uncontrollably within a host but also spread to new hosts via horizontal transmission, and in some cases even vertical transmission (Tissot et al. (2024)). Here, one can envisage both co-evolution and conflict resolution between host and tumour, akin to other internal conflicts such as sex-ratio distortion. Indeed, while vertically transmitted cancers may have been especially important in early multicellular lineages, high prevalence of transmissible cancers in a population is likely to be detrimental in the long run (Aubier et al., 2020) and select for conflict resolution (Hammerschmidt et al., 2014).

In sum, whether cancer is an internal evolutionary conflict is context dependent. With the exception of transmissible cancers, we suggest that cancer generally does not satisfy our criteria for being called an internal evolutionary conflict.

With these considerations in mind, biological interactions can be placed into categories depending on how many of the above criteria they satisfy (Fig. 1). The “internal” party in an internal evolutionary conflict coincides with the notion of a selfish genetic element, as defined by Werren et al. (1988a). However, this alternate conceptualisation allows us to explicitly contrast the study of selfish genetic elements with scenarios that are non-internal evolutionary conflicts such as predator-prey interactions and resource competition (Queller and Strassmann, 2018), internal non-evolutionary conflicts such as cancer (see Box 1), and internal evolutionary non-conflicts such as obligately-vertically transmitted mutualistic endosymbioses with autosomal-like inheritance patterns (Athreya et al., 2025).

Archetypal examples of internal evolutionary conflicts per our definition include those produced by meiotic drivers that distort Mendelian segregation while imposing an organismal-level cost (Lindholm et al., 2016), maternally imprinted genes that control the resources a foetus takes from its mother via the placenta (Barlow et al., 1991), and maternally inherited endosymbionts such as *Wolbachia* that distort organismal sex ratios by feminizing or killing male hosts (Brenninger et al., 2025). We illustrate these various categories in Figure 1.

### Modelling an internal evolutionary conflict

Once we have established that a biological phenomenon is an internal evolutionary conflict, the next step for modelling it requires thinking about how it arises. We propose that there are three potential paths to conflict: conflict due to segregation asymmetry, conflict due to relatedness asymmetry, and conflict due to class-linkage asymmetry. All involve an organism, and a heritable element that is internal to this organism that can spread due to (i) non-Mendelian rules of segregation within each organism (segregation asymmetry), (ii) non-random interactions or mating between organisms (relatedness asymmetry), (iii) non-identical time spent across different classes of organisms (class-linkage asymmetry). In this section, we demonstrate how one can build models of these three paths to conflict. These paths are similar, but not exactly equivalent, to the three types of conflict proposed by Gardner and Úbeda (2017). We explain how the two approaches connect (and why we depart from theirs) in the Discussion, subsection 5.2.1.

We will refer to our focal internal element as the ‘selfish allele’ and the putative conflict is between this element and the organism it resides within. We use ‘selfish’ to refer to situations where the internal element produces a phenotypic effect that pushes the organismal phenotype towards the optimum for that element, but away from the optimal phenotype for the organism at large. Under this definition, selfishness includes cases where some genes benefit from a level of organismal altruism greater than is optimal for the organism (see our section below titled “conflict due to relatedness asymmetry”). Though we use the word ‘allele’ throughout, the physical instantiation of our element could vary from a single nucleotide to an entire chromosome; what matters is that it is stably inherited and segregates as a single unit.

In all models below, we shall assume a diploid-dominant life cycle, non-overlapping generations, and use the phrase ‘well-mixed’ to refer to a population in which all individuals are equally likely to interact with each other. We also neglect spatial structure and environmental fluctuations, and further assume populations are infinitely large. These simplifications are not meant to imply that the phenomena we neglect are unimportant in determining evolutionary outcomes. Though demography (Metcalf and Pavard, 2007), spatial structure (Durrett and Levin, 1994), and finite population effects (Proulx and Day, 2002; Bhat and Guttal, 2025) can all strongly influence and sometimes even reverse evolutionary outcomes, the mathematical techniques required to study the evolutionary consequences of incorporating these phenomena are rather involved (see Beaghton and Burt (2022), Paril and Phillips (2022), and Kläy et al. (2026) for explicit demonstrations in the context of gene drives). Since our goal is to present a widely accessible primer and synthesis that is accessible to those who are new to modelling, we stick to a mathematically friendly framework grounded in classical population genetics. We point keen readers to studies that extend our models of conflict in various biologically relevant directions in a dedicated section towards the end of the manuscript.

In all models, we represent the evolutionary interests of the organism by matching the optimal phenotype from the perspective of the organism with the phenotype produced by a second allele at the same or another locus. The allele representing organismal interests is taken to always be autosomally inherited, whereas some selfish alleles spread due to the fact that they are not autosomally inherited (see section on class-linkage asymmetry). While it is possible to formulate the entire problem as one of differing (inclusive) fitness interests between genes (as done by Gardner and Úbeda, 2017), we choose the above conceptual route since it aligns more closely with our definition of an internal evolutionary conflict. This simplification is also in line with similar conceptual treatments such as the parliament of genes (Leigh, 1977) and the strategic reference gene (Fromhage and Jennions, 2019), which equate the consensus evolutionary interests of a complex genome with those of the organism. By restricting the conflict to one or two loci, we can use simple models that shift the challenge away from the mathematics, instead focusing on making sure the appropriate book-keeping occurs. The goal, then, is to assess whether a selfish allele can persist if it gains an advantage due to segregation, relatedness, or class linkage asymmetry.

Our models will also share a common conceptual progression: in each model, we first consider an allele *E*_0_ that is neutral and represents the organism or parliament of genes. Then, we will consider a putatively selfish allele *E*_*s*_ that is (i) deleterious to the larger organism, but (ii) nevertheless persists in the population by exploiting an asymmetry of some kind. In each case, we will illustrate how organismal interests oppose this selfish allele, by showing that it is purged from the population if it does not exploit the associated asymmetry.

## 4.1 Conflict due to segregation asymmetry

Conflicts due to segregation asymmetry take place when a selfish allele is transmitted more frequently into a heterozygote’s gametes than other alleles at the same locus. Lindholm et al. (2016) review different cases of meiotic drive that can all (at least roughly) be described by the model we introduce below, which is inspired by those compiled in Feldman and Otto (1991). Additional examples include preferentially transmitted B-chromosomes (Jones, 2018) and any homing endonucleases that come with a fitness cost (Burt and Koufopanou, 2004). Early models of transposons (Hickey, 1982) and biased gene conversion (Walsh, 1983) also illustrate their status as examples of conflict due to segregation asymmetry.

### 4.1.1 Constructing a model

Here, we describe a simplified one-locus, two-allele model of meiotic drive, broadly applicable to all of the above mechanisms. Consider a selfish allele *E*_*s*_ and an organism-aligned allele *E*_0_ at a diploid autosomal locus. Let the frequency of *E*_*s*_ be *p* and the frequency of *E*_0_ be 1 − *p. E*_*s*_ affects the proportion of gametes made by an *E*_0_*E*_*s*_ heterozygote, such that *k >* 1*/*2 of them contain *E*_*s*_. This is the transmission distorting effect commonly referred to as drive.

Driving alleles often come with a fitness cost for the organism carrying them (Lindholm et al., 2016). Therefore we will assume that organisms homozygous for the driver allele incur a fitness cost *s* and heterozygotes receive an intermediate penalty modulated by the dominance coefficient *h*. The genotypes *E*_*s*_*E*_*s*_, *E*_*o*_*E*_*s*_, *E*_0_*E*_0_ therefore differ in fitness at the organism level, in the ratio *W* − *s* : *W* − *hs* : *W* (*s >* 0, *h* ∈ (0, 1]).

In a well-mixed population, the frequency of *E*_*s*_ in the next generation is given by

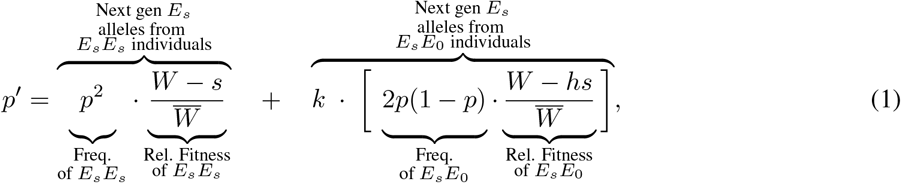

where 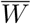 is the mean absolute fitness across the population and is calculated by weighting the fitness of each genotype by its frequency:

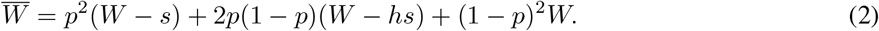

As seen in Eq. 1, dividing a genotype’s absolute fitness by 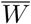 gives its relative fitness in the population – the quantity we need to calculate changes in genotype *frequencies*. The first term on the right hand side (RHS) of Eq. 1 represents the contribution of *E*_*s*_*E*_*s*_ homozygotes towards making *E*_*s*_ alleles in the next generation, whereas the second term represents the contribution of *E*_*s*_*E*_0_ heterozygotes. The drive parameter *k* controls the fraction of *E*_*s*_ gametes made by these *E*_*s*_*E*_0_ heterozygotes. Setting *k* = 0.5 produces Mendelian segregation, whereas when *k >* 0.5 there is transmission distortion.

Now that we have the frequency of the allele in the next generation, we can compute the change in frequency over a single generation, Δ*p*, as

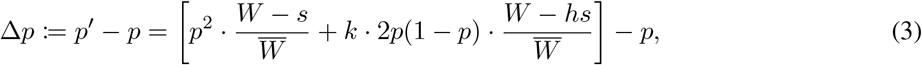

where we have replaced *p*^*′*^ with the RHS of Eq. 1.

Equation 3 can be simplified further, since

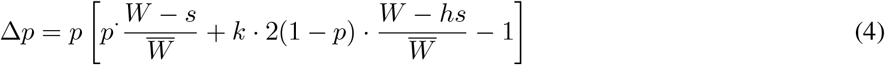

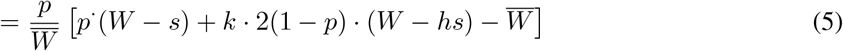

Substituting the definition of 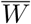 from Eq. 2 within the square brackets now gives

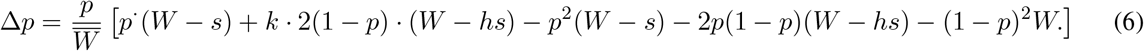

By grouping like terms, we can factor out (1 − *p*) from the RHS of Eq. 6, upon which Eq. 6 takes the form

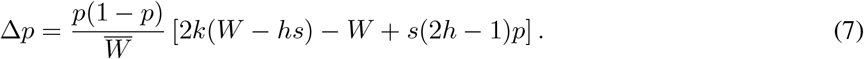

We can now study this model further by asking what value the allele frequency will settle to after many timesteps.

### 4.1.2 Analysing the model

At equilibrium, the frequency of the focal allele must be unchanging from one generation to the next. Thus, we can find potential equilibrium frequencies of the focal allele by setting Δ*p* to zero in Eq. 7. Since the RHS of Eq. 7 is the product of three terms, it can only equal zero if (at least) one of the three terms is individually equal to zero. Δ*p* can thus become equal to zero in three ways, leading to three possible steady states for the frequency of *E*_*s*_: (i) the fixation equilibrium *p* = 1, (ii) the extinction equilibrium *p* = 0, and (iii) the polymorphic equilibrium where *E*_*s*_ is at an intermediate frequency *p*^∗^. To find *p*^∗^, we set Δ*p* = 0 in Eq. 7 and assert that *p* ≠ 0 and *p*≠ 1. This leaves us with 2*k*(*W* − *hs*) − *W* + *s*(2*h* − 1)*p* = 0, which can be rearranged to find that the intermediate equilibrium frequency is given by

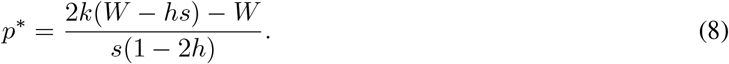

With this knowledge, we are first interested in finding when a selfish allele *E*_*s*_ can invade from rarity. Mathematically rephrased, we are interested in whether an *E*_*s*_ allele introduced at a very small frequency (*p* ≪ 1) will spread (Δ*p >* 0) if the change in frequency is given by Eq. 7. Using either graphical or algebraic criteria (see box 2), one can show that a rare *E*_*s*_ allele will spread if

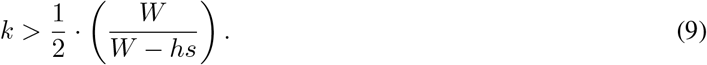

Since *h, s* ≥ 0, the quantity *W* − *hs* can never be greater than *W*, and thus, the fraction *W/*(*W* − *hs*) is at least 1. If the fitness cost associated to *E*_*s*_ is at least slightly visible in the heterozygote (*h >* 0), then *W/*(*W* − *hs*) will be more than 1. Three conclusions can thus be made from inequality 9:

1. in the absence of drive (*k* ≤ 1*/*2), even a slightly deleterious effect at the organismal level (any *s >* 0) will purge the *E*_*s*_ allele from the population. This shows that organismal interests oppose the allele *E*_*s*_ when it does not distort segregation.
2. however, for a given selective disadvantage *s >* 0, *E*_*s*_ can invade when it biases transmission strongly enough towards itself (*k* must be large enough compared to 1*/*2);
3. lastly, if the selective disadvantage *s* is set to zero, i.e. drive is cost-free, then the right hand side of Inequality 9 reduces to 1*/*2, so even the smallest amount of drive is sufficient for invasion;

If the selfish allele *E*_*s*_ spreads when rare, it subsequently has two possible long term fates: the allele could either fix in the population, or remain at an intermediate frequency alongside the non-selfish allele *E*_0_. Using techniques outlined in Box 2, we see that there will be a stable polymorphism between *E*_*s*_ and *E*_0_ if

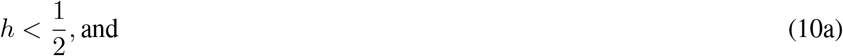

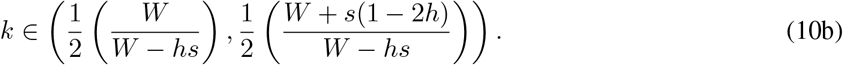

The selfish allele will go to fixation after invading if either of the above inequalities is not satisfied but Ineq. 9 is satisfied. Figure 3 plots the minimum segregation ratio necessary for invasion, for different values of the normalised selective disadvantage *s/W*.

**Box 2: Equilibrium frequencies and evolutionary outcomes**

In this box, we illustrate how one makes conclusions about the invasion and persistence of an allele from models such as ours. In both the segregation asymmetry model (Eq. 7) and the relatedness asymmetry model (Eq. 14), the change in frequency over one generation takes the form

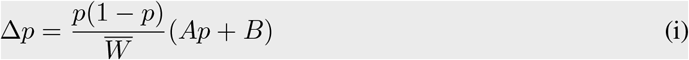

where *A* = *s*(2*h* − 1), *B* = 2*k*(*W* − *hs*) − *W* for the meiotic drive model Eq. 7, and *A* = (*b/*2) − *c*[1 − (*m/*2)], *B* = (1 − *m*) [*c* − (*b/*2)] for the relatedness asymmetry model Eq. 14. The frequency of *E*_*s*_ is at equilibrium when it does not change from generation to generation (by definition). Thus, at an equilibrium, we must have Δ*p* = 0. From the RHS of Eq. i, we see that Δ*p* can equal zero if either:

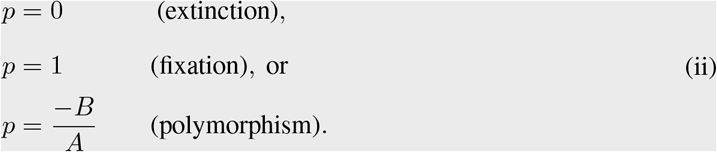

Since *p* is an allele frequency, it must be between 0 and 1, and the polymorphic equilibrium *p* = −*B/A* is thus only biologically meaningful if 0 *<* −*B/A <* 1. If this condition is satisfied, then a polymorphism can exist, whereas if it is not satisfied, then the allele must either reach fixation or go extinct.

Biologically, we are interested in more than whether an allele frequency *p*^∗^ is ‘merely’ an equilibrium for the equation Eq. (i). We also want to check whether a population in which the allele is at a different trait frequency *p*≠ *p*^∗^ evolves to reach the frequency *p*^∗^ over time, a property called *stability* (Otto and Day (2011), chapter 5). For instance, since we do not explicitly incorporate mutational events, an allele frequency of *p* = 0 is always an equilibrium (if there are no copies of a focal allele in a population, its frequency cannot change), but the equilibrium may not be stable (if a copy of the focal allele is introduced into a population that does not initially contain any copies, its frequency may increase).

We can graphically find both equilibria and their stability by plotting Δ*p* against *p*. Intuitively, an equilibrium frequency *p*^∗^ is stable if the allele frequency decreases (Δ*p <* 0) whenever the current frequency is greater than the equilibrium frequency (*p > p*^∗^), and increases (Δ*p >* 0) whenever the current frequency is lower than the equilibrium frequency (*p < p*^∗^). Thus, when plotted, an equilibrium frequency *p*^∗^ is stable if the curve defined by Eq. i is above the *x*-axis whenever we are to the left of *p*^∗^ and below the *x*-axis whenever we are to the right of *p*^∗^. Without going into further detail, you can use this criterion to work out all possible behaviours of the system either graphically or algebraically. The mathematical translation of the above statements is in terms of the derivative of Δ*p* with respect to *p*, which can be used to measure if Δ*p* is increasing or decreasing at the equilibrium point. In particular, an equilibrium frequency *p*^∗^ is stable if

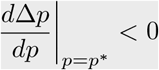

where the big line on the right denotes that the derivative must be taken and then evaluated at *p*^∗^ when one wishes to use this criterion.

The final outcome is that there are four possible behaviours of the system based on the signs and relative magnitudes of *A* and *B*, which we illustrate in Fig. 2. We encourage interested readers to experiment with *A* and *B* in Eq. i to replicate the results summarised in Fig 2 as a useful exercise, or also by computing the derivatives required above. Chapter 5 of Otto and Day (2011) provides a more detailed pedagogical introduction to equilibria and stability of dynamical systems such as Eq. i.

**Figure 2.**
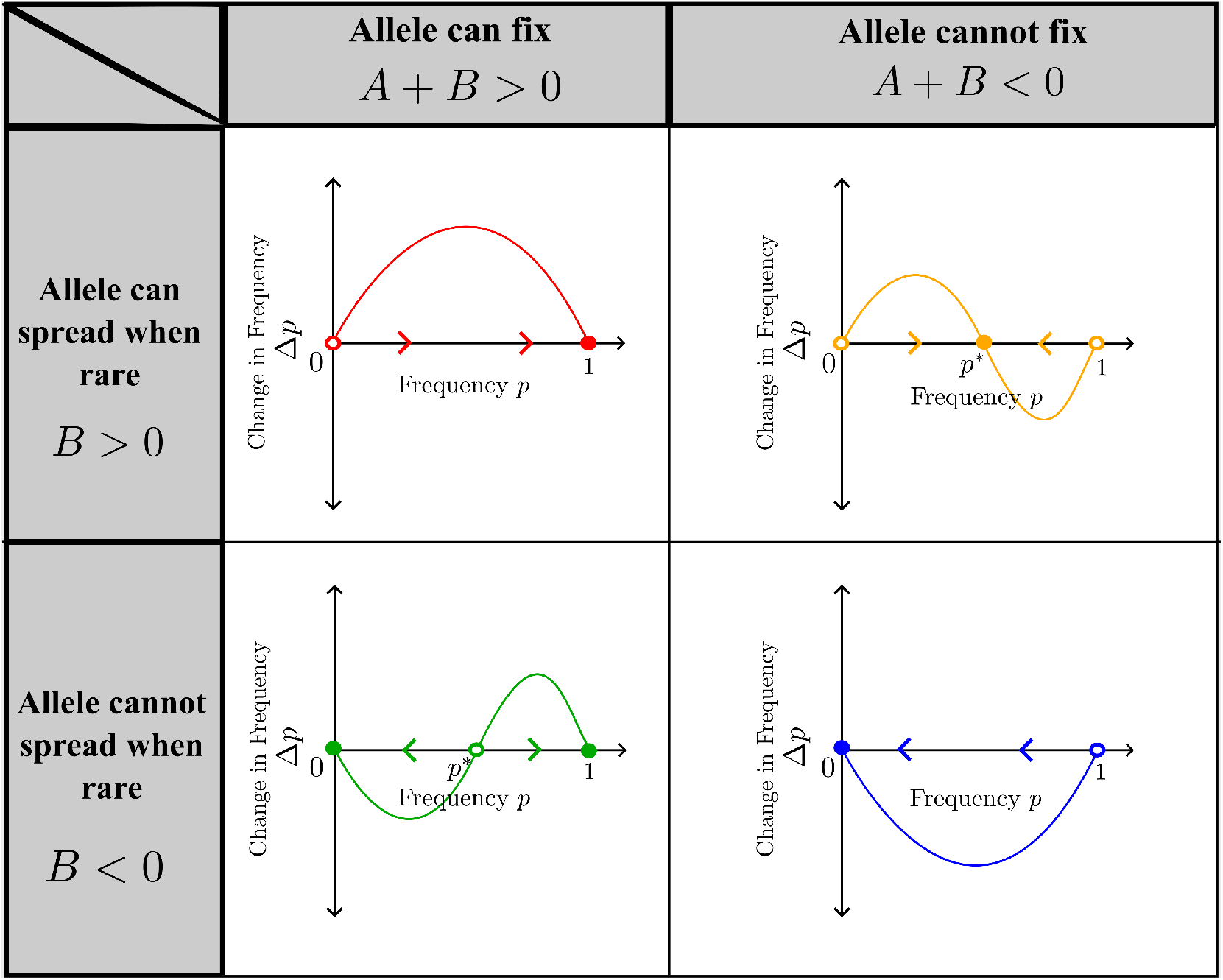
All possible dynamical behaviours of the system Δ*p* = *p*(1 − *p*)(*Ap* +*B*). Points at which Δ*p* = 0 (i.e. the curve intersects the *x*-axis) are equilibria. Arrows indicate the direction of evolution (change in trait frequency) when the population has not yet reached equilibrium. Filled circles indicate that the equilibrium is *stable*, i.e. that populations close to this equilibrium will evolve towards it. Empty circles indicate that the equilibrium is unstable. Since *p* is a frequency, existence of the polymorphic equilibrium (top right and bottom left) also requires − *B/A* to be between 0 and 1. We are interested in the top row, where a rare allele is able to spread/invade. **Alt text**: Four panels indicating what the graph of Δ*p* vs. *p* must resemble for each of the dynamical behaviours to take place. The figure shows that an allele can invade from rarity if and only if this graph is above zero for small values of *p*.

In sum, segregation-distorting alleles can persist even if they impart an organism-level selective disadvantage, as long as the advantage they derive from being over-represented at segregation is high enough. For studying segregation distortion in other systems, this approach can be expanded by adding biological details relevant to the system and question. In SI section S7, we show how this might be accomplished by extending the model above to sex-specific distortion, homozygote lethality and multiple mating with sperm competition.

Many models of, and arguments around hierarchical evolutionary processes make use of the Price equation (Price, 1970; Price, 1972; Frank, 2012; Gardner, 2020), including to formalise notions of internal evolutionary conflict (Okasha, 2006; Patten et al., 2023). Beloved for its generality, it is mathematically equivalent to population genetic models as above, and can be extended to include relevant complexities such as kin-selection (Taylor, 1990b; Taylor and Frank, 1996; Queller, 2017). In SI section S8, we mirror the above analysis using the Price equation to demonstrate this equivalence.

## 4.2 Conflict due to relatedness asymmetry

The next grouping of selfish alleles that fall within our classification of internal evolutionary conflicts are those that exploit relatedness asymmetries. Relatedness here refers to the probability that a randomly chosen gamete of a social partner carries a common allele with the focal individual, from the perspective of the focal, selfish locus. These alleles need not be identical by descent. The conflict manifests through some biasing of which social partners a fitness-affecting behaviour is directed towards, such that relatedness at the focal locus differs from relatedness at other, unlinked loci. This assortment asymmetry can then lead the expression of a certain social behaviour to be deleterious for the majority of the genes within an organism, but beneficial for alleles at the focal locus.

An evocative example of conflict due to relatedness asymmetry can manifest when alleles produce costly ‘greenbeard’ phenotypes. Costly greenbeard alleles are those that code for costly altruism (non-costly greenbeards can exist, Gardner and West, 2010a, but do not cause conflict), and simultaneously produce a signal that carriers of this allele use to preferentially direct cooperative behaviour exclusively to other carriers of the allele (Hamilton, 1964a; Dawkins, 1976; Gardner and West, 2010b; Madgwick et al., 2019). Greenbeards do not cause internal evolutionary conflict in many cases (Ridley and Grafen, 1981; Gardner and West, 2010a; Biernaskie et al., 2011; Aubier and Lerch, 2024), except if genealogical relatives interact preferentially with each other and a costly greenbeard allele arises that restricts altruism only to those genealogical relatives that also possess a greenbeard (Biernaskie et al., 2011). In particular, greenbeard alleles can be in conflict with alleles that code for beard-indiscriminate helping, since assortment between greenbeards is not guaranteed by assortment between genealogical relatives. This relatedness asymmetry implies that the kin-selection benefit of helping an unbearded sibling is lower for the greenbeard gene than for the indiscriminate helping gene.

While the greenbeard example is useful for explaining relatedness asymmetries, it is not well supported empirically. We instead choose to model genomic imprinting, which requires relatedness asymmetries of the same kind. Genomic imprinting is parent-of-origin-dependent expression of an allele, resulting in a trait value that depends on which parent – mother or father – the allele was inherited from. The kinship theory of genomic imprinting (Haig, 2000) is built on the premise that with random mating and polyandry, there is no statistical association between a paternally derived allele and the allele copies present in an individual’s mother. This means that although there is strong genealogical relatedness between mother and all her offspring, relatedness specific to paternally derived alleles among these maternal siblings is zero. Imprinted alleles inherited paternally should therefore produce phenotypes that are less altruistic towards the organism’s mother (and her future offspring) than would be expected in the absence of imprinting. By contrast, if the same allele is inherited maternally, gametes produced by mother and offspring have a high probability of carrying like allele copies. The future reproductive output of the focal organism’s mother is thus of drastically different importance depending on an allele’s parent of origin, generating the potential for internal evolutionary conflict.

Spurred on by the discovery of imprinted genes that control growth in the womb (Barlow et al., 1991; DeChiara et al., 1991), the first models of imprinting were constructed to explain how much a focal offspring is selected to “take” from its mother during development (e.g. Mochizuki et al., 1996; Haig, 1997; Van Cleve et al., 2010). Here we present a population genetic caricature of these models that focuses instead on altruism between maternal siblings, inspired by Supplementary Note 4 in Scott and West (2019). Altruism is analogous to the womb growth idea above since selfishly taking “too much” from one’s mother is similar to being selfish with respect to one’s maternal siblings, who must also grow in the same mother’s womb.

### 4.2.1 Constructing a model

Consider alleles *E*_*s*_ and *E*_0_ at a locus that affects altruism during social interactions. Expression of the *E*_*s*_ allele comes at a personal cost *c* to the organism performing the altruistic behaviour, but produces a benefit *b* that is conferred to their social partner. We assume that the altruism allele is completely dominant, such that in *E*_*s*_*E*_0_ heterozygotes, the personal cost incurred and benefit conferred to social partner are equal to that in *E*_*s*_*E*_*s*_ homozygotes (for incomplete dominance, see Supplementary Note 4 in (Scott and West, 2019)). The allele *E*_0_ is effectively neutral; it does not code for altruism and is not imprinted, whereas the allele *E*_*s*_ coding for costly altruism is silenced with probability *m* when inherited paternally. To get a feel for how this works, consider the case where *m* = 0: there is no effect of imprinting, and *E*_*s*_ is expressed identically regardless of its parent-of-origin. In contrast, when *m* = 1, *E*_*s*_ is only expressed when maternally inherited; the strongest possible case of imprinting in our model. With this model, our goals are to i) derive the evolutionary fate of the *E*_*s*_ allele invading from rarity and ii) assess how systematically varying *m* changes this fate.

Adult diploids are produced following random mating, and fitness is affected by their social interactions. We impose that these interactions occur exclusively between individuals that share a maternally inherited allele at our focal locus; interactions with respect to the paternal allele are taken to be well-mixed. Note that we are thus assuming that mating is random but that social interactions are non-random. These are clearly strong assumptions, but we make them to best illustrate the effect of imprinting. Our model is therefore an informative toy, though weaker forms of such assortment-biases between maternal siblings may arise due to sex-biased dispersal patterns, or if many organisms have the same mother but most have different fathers (see e.g. Van Cleve et al., 2010). Different demographic regimes could also give rise to paternal assortment biases, for example in sex-role reversed species where males care for clutches they have sired with multiple females (e.g. sticklebacks; Barbasch et al., 2026). Our model can easily be extended in that direction by flipping the interaction structure, though that is beyond the scope of this manuscript.

Let the current frequency of *E*_*s*_ be *p*. Four genotypes exist in this population: (i) *E*_*s*_*E*_*s*_ homozygotes, which always inherit an *E*_*s*_ from their mother, (ii) *E*_*s*_*E*_0_ heterozygotes that inherit a maternally derived *E*_*s*_, (iii) the remaining half of heterozygotes that inherit a *E*_*s*_ allele from their father, and (iv) *E*_0_*E*_0_ homozygotes that don’t carry the *E*_*s*_ allele. For clarity, we list the maternally derived allele first when writing the genotypes, such that *E*_*s*_*E*_0_ refers to organisms that inherited a *E*_*s*_ from their mother and a *E*_0_ from their father. Among these, (i) and (ii) will always express the altruistic phenotype, organisms of (iii) express it with probability 1 − *m*, and (iv) will never do so. The fitness accrued by each of these classes is different, and found by writing down the types of interactions that lead to a fitness effect, how frequently each type of interaction takes place, and how much this effect amounts to. The cost of altruism is always personally incurred, and uniformly applied against the base fitness of 1, and only depends on whether the focal organism expresses the allele *E*_*s*_. In other words, the absolute fitness of every genotype is of the form 1 + (effect of social interactions), where the effect of social interactions can be read off from Table 1. For instance, the absolute fitness of *E*_0_*E*_*s*_ organisms (the single genotype where paternal silencing of *E*_*s*_ is possible in our model) is computed as

**Table 1.** All pathways of *E*_*s*_ changing in frequency. Genotypes are written with the maternally inherited allele first, e.g. the genotype *E*_*s*_*E*_0_ denotes an organism which inherited *E*_*s*_ from its mother and *E*_0_ from its father. We assume the baseline fitness, independent of social interactions, is identically 1 for all genotypes, and exclude it from this table.

| Focal genotype | Genotypic frequency | Social partner | Probability of encounter | Fitness from social interaction |
| --- | --- | --- | --- | --- |
| $E_s E_s$ | $p^2$ | $E_s E_s$ | $p$ | $b - c$ |
| | | $E_s E_0$ | $1 - p$ | $b - c$ |
| $E_s E_0$ | $p(1 - p)$ | $E_s E_s$ | $p$ | $b - c$ |
| | | $E_s E_0$ | $1 - p$ | $b - c$ |
| $E_0 E_s$ | $(1 - p)p$ | $E_0 E_s$ | $p$ | $m \cdot 0 + (1 - m) \cdot (b - c)$ |
| | | $E_0 E_0$ | $1 - p$ | $m \cdot 0 + (1 - m) \cdot (-c)$ |

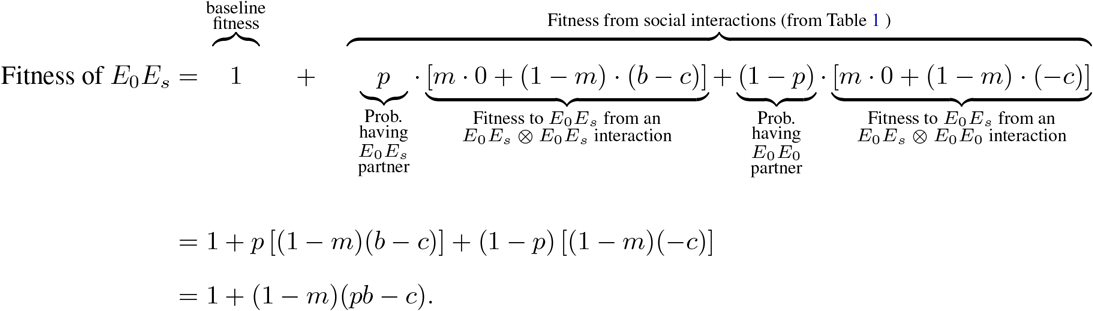

After repeating this process for all relevant genotypes, we can compute the frequency of *E*_*s*_ alleles in the next generation as

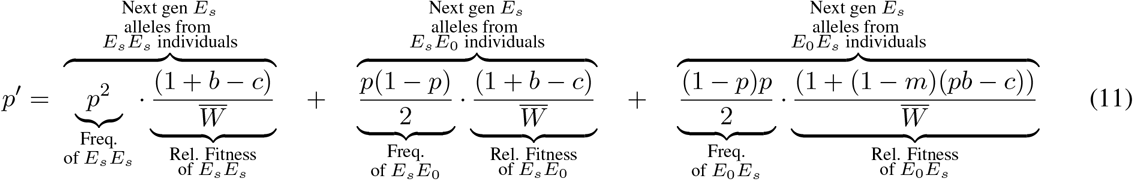

where 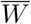 is the mean fitness of the population, given by

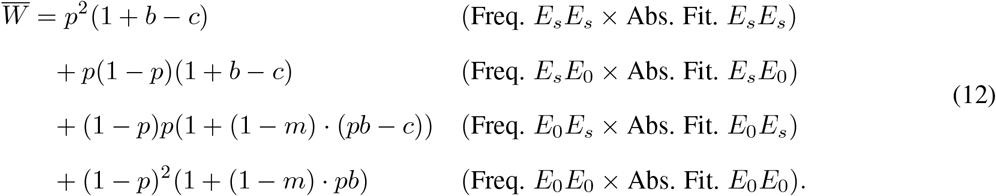

As before, we can now compute the change in frequency over a single generation, Δ*p* := *p*^*′*^ − *p*, as

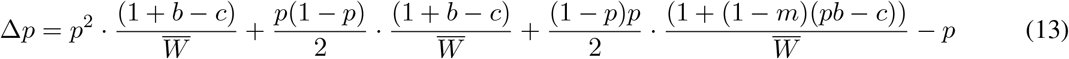

which can now be simplified further and analysed systematically.

### 4.2.2 Analysing the model

Grouping and rearranging the terms on the RHS of Eq. 13, we can rewrite Eq. 13 as

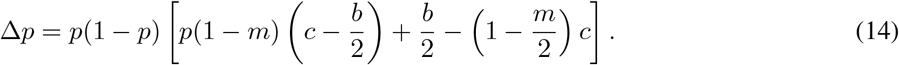

Eq. 14 predicts three potential long-term outcomes for *E*_*s*_ (see box 2): fixation, extinction, or polymorphism where *E*_*s*_ coexists with the other allele at frequency

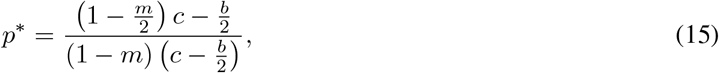

To understand organismal interests, consider an un-imprinted version of this allele (*m* = 0). For such a case, the only outcomes are extinction or fixation since the internal equilibrium (15) coincides with the fixation equilibrium *p* = 1. The conditions in box 2 reveal the invasion criterion: if 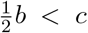, the allele *E*_*s*_ is purged from the population, and if 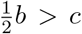, it goes to fixation. If these inequalities are reminiscent of Hamilton’s rule, *rb > c* (Hamilton, 1964b), it is because they are indeed versions of Hamilton’s rule. The 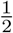 in our inequality is the relatedness coefficient *r*, which we arrive at because social partners always have the same maternally-inherited allele, whereas interactions are random with respect to the paternally derived allele. The probability that a gamete produced by both interactants carries the same allele is therefore 0.5 averaged across many interactions. Invasion always implies fixation for this un-imprinted allele since there is no negative frequency-dependent selection – as the frequency of *E*_*s*_-bearing altruists increases, they meet each other only more often. Therefore, under this pattern of assortment by maternally-inherited allele, an unimprinted *E*_*s*_ will invade if and only if *b >* 2*c*, and will otherwise be purged by selection.

Now consider cases where *E*_*s*_ can be paternally silenced. As soon as *m >* 0, the conditions under which *E*_*s*_ can invade (the equilibrium *p* = 0 must be unstable, see Box 2) expands from *b >* 2*c* to

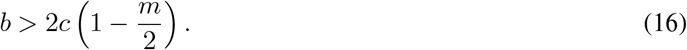

Specifically, invasion is now possible for the additional values of *b* between 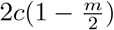 and 2*c*; conditions under which an unimprinted analogue would be purged by selection. This is because imprinting prevents *E*_0_*E*_*s*_ heterozygotes with paternally-inherited *E*_*s*_ from “leaking” the benefits of the social behaviour to *E*_0_*E*_0_ homozygotes (which do not carry *E*_*s*_ and are analogous in this setting to cheaters, see Fig. 4a). Unlinked genes at other loci always bear the (organism-wide) cost of the altruistic behaviour, but do not benefit from the reduced leakage. This is because interactions take place between organisms who share a maternally inherited allele at the imprinted locus, whereas the probability that the interaction partner is identical by state at these other loci is indistinguishable from if the population was well-mixed (interacting at random). In other words, relatedness at these other loci is zero. Genomic imprinting thus constitutes an internal evolutionary conflict, since it allows the persistence of an allele that is opposed by organismal interests. Figure 4b shows the minimum imprinting strength required for the invasion of *E*_*s*_ for different cost-benefit ratios *c/b*.

**Figure 3.**
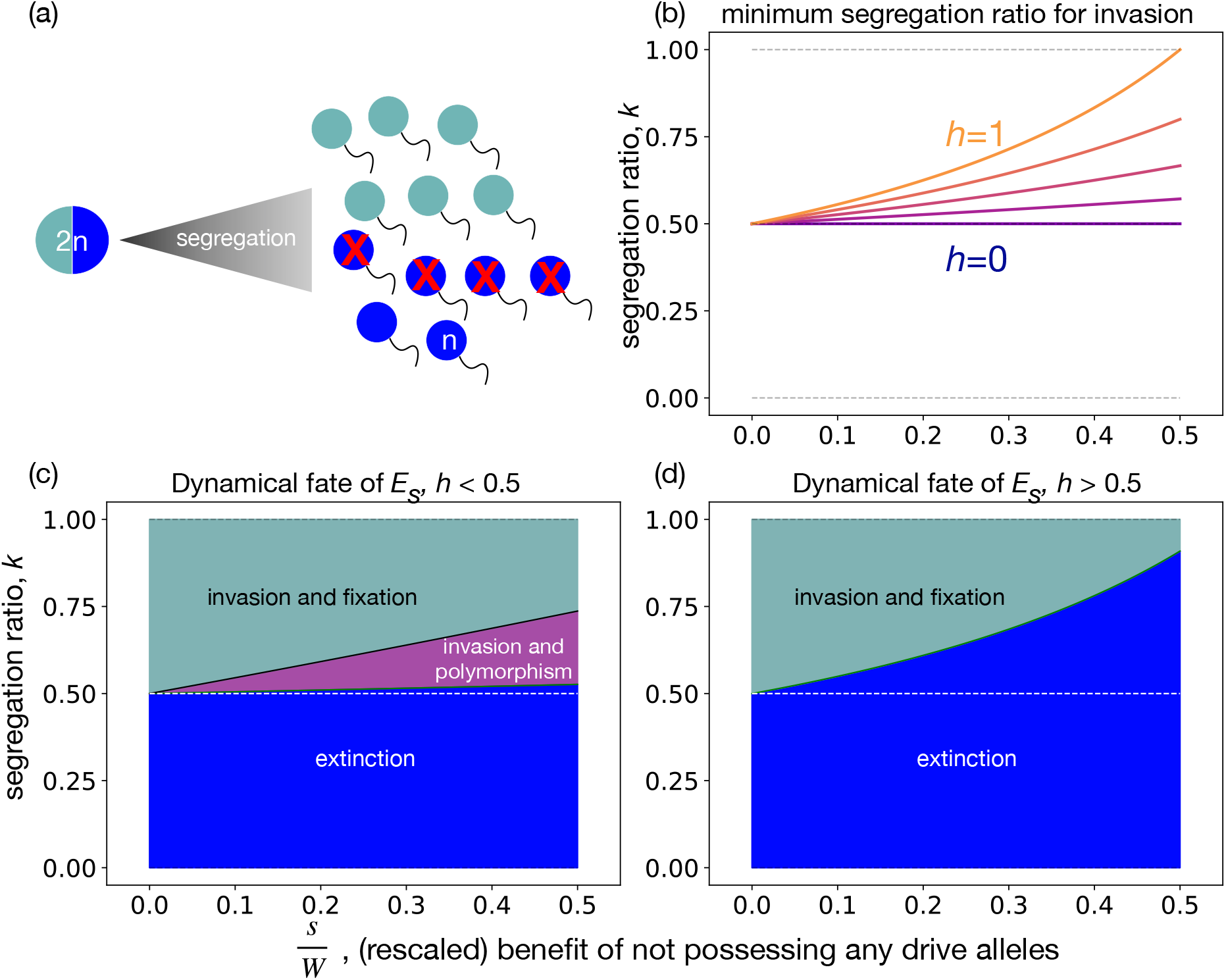
Invasion and persistence of costly segregation distortion. **(a)** One mechanism by which conflict due to segregation asymmetry may occur. This visual is modelled after the *Drosophila* Segregation Distorter system, where the allele analogous to our *E*_*s*_ produces a diffusing toxin and a private antidote to this toxin, which has the effect of killing un-like gametes that carry the *E*_0_ allele. **(b)**The minimum segregation ratio required for invasion of a deleterious allele, as a function of its fitness cost (Eq. 9). Invasion of an allele with higher fitness cost requires a stronger distortion of segregation in the allele’s favour; the required strength increases with the dominance of the deleterious allele. Given that *k* is high enough for invasion, both fixation and stable polymorphism are possible (see 10b): when *h <* 0.5 (panel **c**, *h* = 0.1), whereas an invading allele will always go to fixation if *h >* 0.5 (panel *d, h* = 0.9). **Alt text**: Four panels. (a) shows a half blue-half grey heterozygote producing blue and grey gametes, blue gametes have a red X marked on them. Next three panels plot segregation ratio *k vs. s/W*, the (rescaled) fitness benefit of not possessing any drive alleles. Panel (b) shows multiple lines, for different values of *h*. All are increasing, and never below *k* = 1*/*2. The line for *h* = 1 is flat at *k* = 1*/*2, and the line for *h* = 0 is farthest from *k* = 1*/*2. Panel (c) plots two lines, never below *k* = 1*/*2. Area below both lines is labelled “extinction”, area above both labelled “invasion and polymorphism”, area between lines is labelled “invasion and fixation”. Panel (d) plots one line, never below *k* = 1*/*2. Area below is labelled “extinction”, area above labelled “invasion and fixation”.

**Figure 4.**
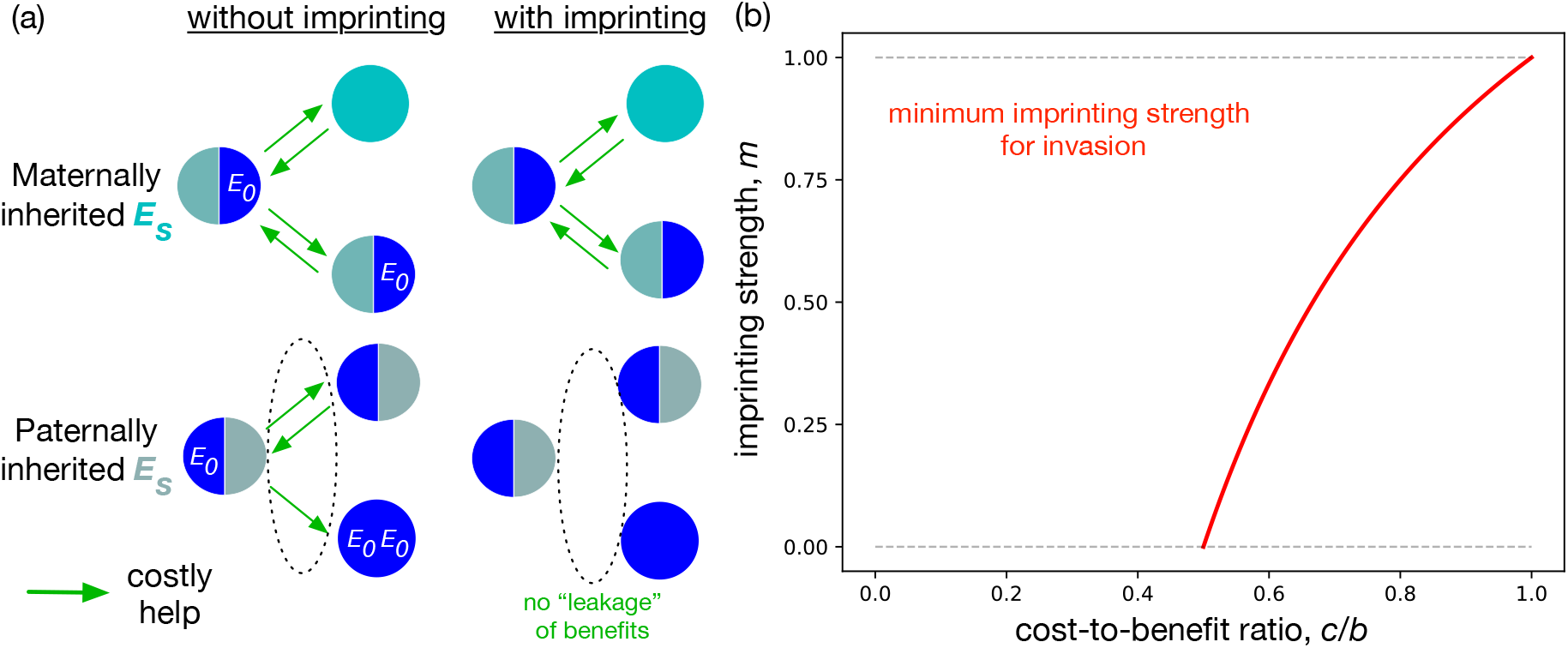
Invasion of (maternal) genomic imprinting. (a) A graphical illustration of why maternal imprinting is beneficial under maternal sibling-biased interactions as we have modelled them. Under maternal imprinting, heterozygotes that inherited the altruism allele from their father stop helping (bottom right) the non-altruistic *E*_0_*E*_0_ homozygotes that they would have helped (bottom left), had they not been maternally imprinted. The latter class of individuals are cheaters, since they reap the benefit of costly helping by another, but do not help in return. (b) For cost-to-benefit ratio *c/b <* 0.5, imprinting is unnecessary, i.e an unimprinted allele can invade into the population. However, as *c/b* increases above 0.5, imprinting is necessary for invasion, and the necessary strength increases with the deviation of *c/b* compared to 0.5. This deviation is the correct analogue to the deleterious fitness disadvantage *s* analysed in the previous section and depicted in Fig. 3. Segregation is assumed Mendelian, no other parameters. Invasion always implies fixation in this model; in other words, there is no negative frequency-dependent selection on the allele *E*_*s*_. **Alt text**: Two panels. (a) depicts half-blue half-grey heterozygotes helping (and being helped) by those they share a maternally-inherited allele with. Panel (b) is a plot showing a line that is non-zero only for cost-benefit-ratio above 0.5, and increases smoothly to the value 1 when cost-benefit-ratio equals 1.

While an un-imprinted allele with *m* = 0 that invades always goes to fixation, an imprinted allele can potentially settle at a polymorphic equilibrium that involves coexistence with the other allele *E*_0_. This takes place when the invasion criterion (16) is satisfied, and simultaneously the polymorphic equilibrium must be dynamically stable. The methods of Box 2 show that this outcome takes place if and only if *b < c*, which is a condition that would prevent invasion in the first place (see (16): if invasion must take place, the lowest *b* can get is to be exactly equal to *c*, which takes place when *m* = 1). Therefore, a polymorphic equilibrium is never stable under this model, and imprinted alleles that invade also go to fixation.

## 4.3 Conflict due to class-linkage asymmetry

The third form of conflict that qualifies under our definition takes place between two genomic regions that have different modes of inheritance in different classes of organisms. Consider, for example, the contrast in inheritance between autosomes and the matrilineally inherited *Wolbachia* endosymbiont that infects arthropod taxa (Werren, 1997). Let an autosomal gene represent the evolutionary interests of the organism as a whole and take the sexes as the relevant classes. *Wolbachia* can only be transmitted by females and are therefore selected to optimise matrilineal transmission, whereas autosomes – transmitted in equal proportions by females and males – are selected to optimise transmission through both sexes. This discrepancy over the *value* of the male transmission pathway produces the potential for internal evolutionary conflict, which is exemplified by the widespread expression of sex-allocation distortion, male-killing and feminisation in *Wolbachia* infected host species (Brenninger et al., 2025).

The value of a given class for allele propagation is the key to modelling this third form of internal evolutionary conflict. Formally, this concept is captured by the notion of juvenile class reproductive values (Fisher, 1930; Taylor, 1990b; Rautiala and Lehtonen, 2024). Loosely defined, the juvenile class reproductive value is the probability that a randomly sampled allele present many generations in the future, is currently found within an organism belonging to the specified class (for an explanation for why we specify juvenile cohorts see Grafen, 2014). In our *Wolbachia* example, males never transmit alleles and thus the total reproductive value of juvenile males is zero. The future allele therefore must have resided in a contemporary female, yielding a total reproductive value of current juvenile females equal to one. If the allele is instead autosomal, the fact that all organisms must receive one autosomal allele copy from their mother and another from their father (in diplo-diploids) means that the future allele descends from a juvenile male or female with equal probability. The above line of reasoning leads us to the notion of class linkage.

An allele can be said to be class-linked if its class-specific reproductive value is “unusually” concentrated in a specific class (more than 1*/N*, for *N* possible classes). Again taking the sexes as the relevant classes, autosomes by definition are never sex-linked, however other alleles might be: an asymmetry in class linkage pattern between two alleles implies a difference in how much fitness effects acting on organisms of each class (females, males) matter for allele propagation. Consequently, an allele coding for some phenotype might be purged by selection if it was present on an autosome, but not if it was found within a matrilineally inherited endosymbiont, thereby producing a conflict. In SI S10 we demonstrate, in a simple setting, how class reproductive values affect invasion criteria.

The complete lack of value males offer (Keaney et al., 2020) to *Wolbachia* allele propagation presents a clear illustration of class-linkage conflict, but removes important nuances that often emerge when modelling sex-specific transmission modes. We therefore focus our investigation on a subtler, but potentially very common, conflict of this kind between alleles on autosomes and X-chromosomes over the expression of sexually antagonistic phenotypes. Specifically, we refer to alleles affecting traits that are expressed by both sexes, with one allele that moves the expressed phenotype closer to the female phenotypic fitness optimum, and a second that moves the phenotype closer to the male optimum. The result is intralocus sexual conflict, where selection is antagonistic, and the sign of selection affecting a given allele depends on the class they are currently expressed in (Schenkel et al., 2018). Note that the internal evolutionary conflict is not that between males and females over expression of the trait (above, we explicitly disqualified intralocus sexual conflict from our definition). Instead, it is between a neutral autosomal allele, which we use as a representation of the organism in aggregate, and any X-linked allele affecting this trait. Below, we construct a mathematical model of this scenario, and show that autosomes and X-chromosomes have different class linkage patterns and therefore “withstand” different levels of sexual antagonism (Frank and Crespi, 2011). However, such conflicts are not limited to X-linked and autosomal genes, and can also take place for example between any combination of genes found on autosomes, X chromosomes and Y-chromosomes (Ågren et al., 2019).

### 4.3.1 Constructing a model

Consider a trait present in a population with separate sexes and XY sex chromosomes, that is affected by allelic variation, should it be present, at several loci. Both sexes express the trait, but the trait value associated with the highest fitness is different for females and males. Consider two alleles *A*_*s*_ and *X*_*s*_ that move the trait value towards the optimum phenotype in one of the sexes, located on an autosome and X chromosome, respectively. Both alleles produce identical phenotypic effects, with a fitness advantage *s*_f_ is females, and *s*_m_ in males, modulated by a dominance coefficient *h* whenever heterozygous (see Table 2 for a full description of the fitness scheme).

**Table 2.** Fitness scheme for a sexually antagonistic allele, when present on an autosome (top row-group), or on an X-chromosome (bottom row-group).

|  |  | Females |  |  | Males |  |  |
| --- | --- | --- | --- | --- | --- | --- | --- |
| Autosomal, $A_s$ | Genotype | $A_s A_s$ | $A_s A_0$ | $A_0 A_0$ | $A_s A_s$ | $A_s A_0$ | $A_0 A_0$ |
| | Fitness | $1 + s_f$ | $1 + h s_f$ | 1 | $1 + s_m$ | $1 + h s_m$ | 1 |
| X-linked, $X_s$ | Genotype | $X_s X_s$ | $X_s X_0$ | $X_0 X_0$ | $X_s Y$ | $X_0 Y$ | |
| | Fitness | $1 + s_f$ | $1 + h s_f$ | 1 | $1 + s_m$ | 1 | |

Heterozygous genotypes are present in both sexes at the autosomal locus, whereas they only occur in females at the X-linked locus. We assume that there is no recombination between the X and the Y, and that the corresponding allele on the Y chromosome has no effect on the sexually antagonistic trait. To maintain a like comparison between the X and the autosomes, we assume that dominance is the same at both loci.

While we have set this up as a two-locus phenomenon, here we will not track co-evolution across chromosomes. Instead, we focus on illustrating the existence of parameter regimes where sexually antagonistic alleles can invade when governed by one rule of inheritance but not by the other. That is, if there was no allelic variation at the X-linked locus, when would *A*_*s*_ invade at the autosomal locus, and vice versa? The discrepancy between the two scenarios reveals the conditions under which there is an internal evolutionary conflict.

With fitnesses defined, we next move to calculating allele frequency changes from one generation to the next. Several changes to our usual approach are required. First, we must build separate recurrences for males and females, since the alleles have sex-specific fitness effects. To show that sex-specific selection is changing allele frequencies within the sexes, we also switch to tracking allele frequencies within the gametes of breeding adults. If we were instead to track allele frequencies amongst zygotes, we would never observe this effect (despite it occurring) for autosomal loci, because the equal contribution of the sexes to both sons and daughters equalises allele frequencies across the sexes (at the population level). Let *p*_A,f_, *p*_A,m_ denote the frequency of *A*_*s*_ at the autosomal locus amongst the female and male gamete pools, respectively. If the locus is instead X-linked, let *p*_X,f_, *p*_X,m_ be the frequency of *X*_*s*_ in these gametes. With an even primary sex ratio, the absence of parent-of-origin effects (i.e. no genomic imprinting) and fair meiosis, the frequency of *A*_*s*_ in the next generation is given respectively by

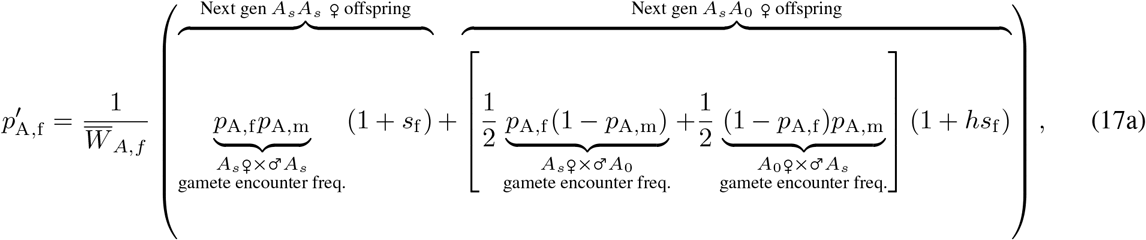

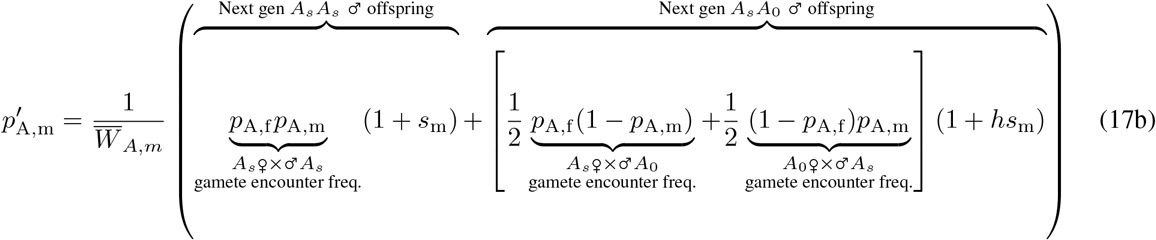

where the sexes (♂*/*♀) in the underbraces indicate the parent the allele originated from (paternal/maternal respectively). The mean fitness in females and males is respectively

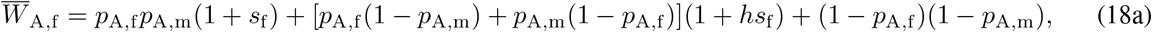

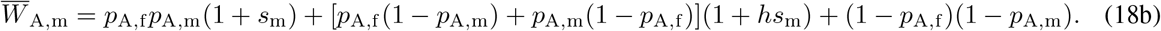

In contrast, the frequency of *X*_*s*_ in the next generation is given in females and males respectively by

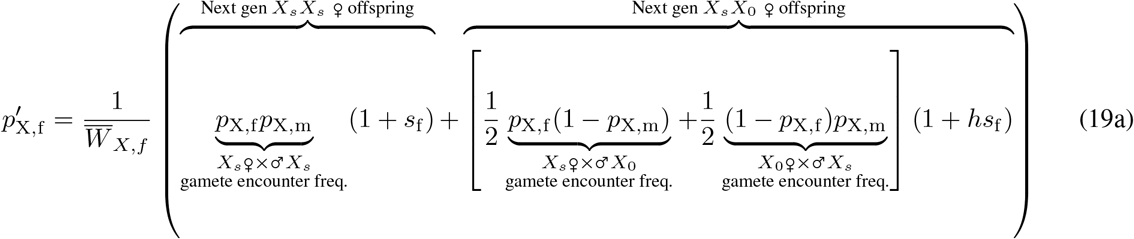

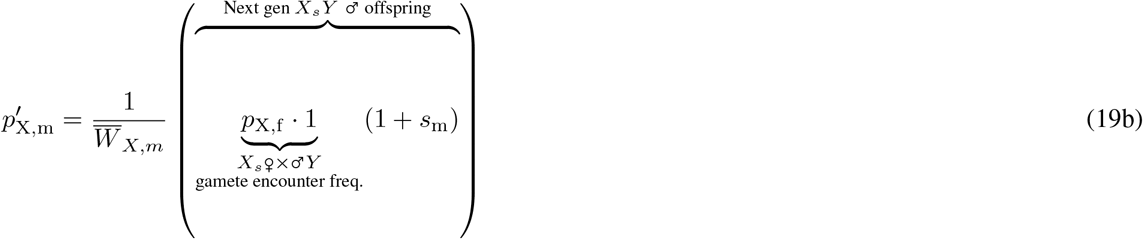

with mean fitnesses

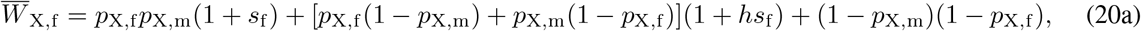

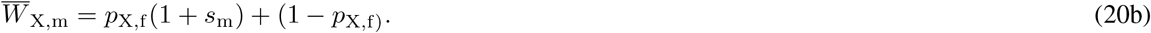

Note that Eq. 19b multiplies the frequency of the *X*_*s*_ allele present in breeding females with its fitness when expressed in males. This is because males inherit their X chromosome from their mother; the frequency amongst male offspring is therefore the frequency of *X*_*s*_ across all eggs produced by the previous generations breeding females. The lifecycle is completed by moving from male offspring to breeding males, which is done by multiplying their frequency by their fitness.

### 4.3.2 Analysing the model

To understand when a given allele – either *A*_*s*_ or *X*_*s*_ – can invade, we must again check whether a small amount of that allele can increase from rarity. However, checking local (in)stability for equilibria of a system of 2 recurrence equations requires slightly more complicated tools than we have so far encountered. In particular, because these systems involve simultaneously tracking the change of two quantities that depend on each other, finding the equilibrium values of the system and checking for its linear stability requires a tool from multivariate calculus and matrix algebra called the *Jacobian matrix*. We have included a short introduction to Jacobians and eigenvalue analysis in the SI (§ S9), and also mention how it connects to the simpler methods in Box 2. In the main text, we focus instead on the final result of using these methods.

To enforce sexually antagonistic fitness effects, we set *s*_m_ = *M, s*_f_ = −*γM*, such that *M* controls the size of the fitness effect changing the trait value has, and *γ >* 0 scales the relative magnitude of the fitness effect in females. For *M >* 0, the behaviour is male-beneficial (and female-deleterious), when *M <* 0, it is instead female-beneficial (following Frank and Patten (2020)). Parsons (1961) showed (with different notation), that *A*_*s*_ can invade at the autosomal locus when

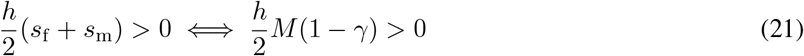

that is, a sexually antagonistic allele at an autosomal locus can invade when the benefit it provides to one sex is larger than the cost it provides to the other; it should just be on average beneficial. Since the dominance *h* is never negative, it can never qualitatively affect this invasion outcome. When *A*_*s*_ is female-beneficial (*M <* 0), the scaling factor *γ* must be larger than one for invasion, and when the allele is female-deleterious (*M >* 0), *γ* must be smaller than one. Therefore, organismal, i.e. autosomal, interests oppose this allele whenever *M* (1 − *γ*) *<* 0.

Can an allele coding for the same deleterious fitness effects persist if it has different class linkage, in particular if it is X-linked like *X*_*s*_? The invasion conditions are indeed different, and given by^1^

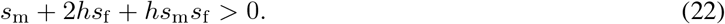

When *X*_*s*_ is male-beneficial and female-deleterious (*M >* 0), the criterion above reduces to

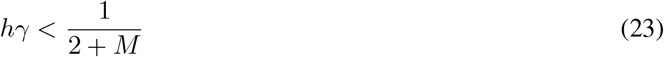

If *X*_*s*_ is instead female-beneficial and male-deleterious (*M <* 0), then it invades from rarity when

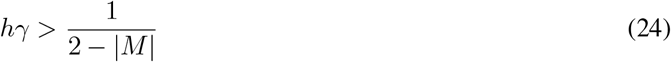

Conflict between *X*_*s*_ and the organism-representing *A*_0_ arises because extinction of a deleterious autosomal allele *A*_*s*_ (*M* (1 − *γ*) *<* 0) does not always imply extinction of the analogously deleterious X-chromosomal allele *X*_*s*_. To see the conflict more clearly, fix the value of *M* and then ask which values of *γ* lead to invasion for the Xchromosomal allele under conditions where *A*_*s*_ would *not* invade. When *M >* 0, the autosomal allele cannot invade when *γ* ≥ 1. The X-chromosomal allele, on the other hand, can invade when

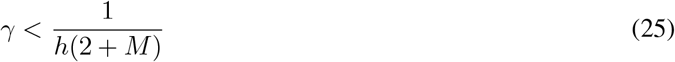

which is ≥ 1 whenever the dominance *h* ≤ 1*/*(2 + *M*). Since we started with *M >* 0, the above criterion implies that the allele must be at least partially recessive.

Alternatively when *M <* 0, the autosomal allele cannot invade anytime *γ* ≤ 1. The X-chromosomal allele, on the other hand, can invade when

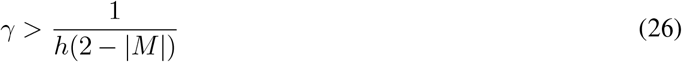

which is ≤ 1 whenever the dominance *h* ≥ 1*/*(2 − |*M* |), which via *M <* 0 implies that the allele must be at least partially dominant.

Therefore, over time, the X chromosome should become enriched for recessive female-deleterious, and dominant female-beneficial alleles that are too costly to spread on an autosome (see Frank and Patten, 2020, for a more genetically explicit version of this model). In Figure 5, we illustrate graphically when both alleles *A*_*s*_, *X*_*s*_ would invade, when neither would, and when there is a conflict.

**Figure 5.**
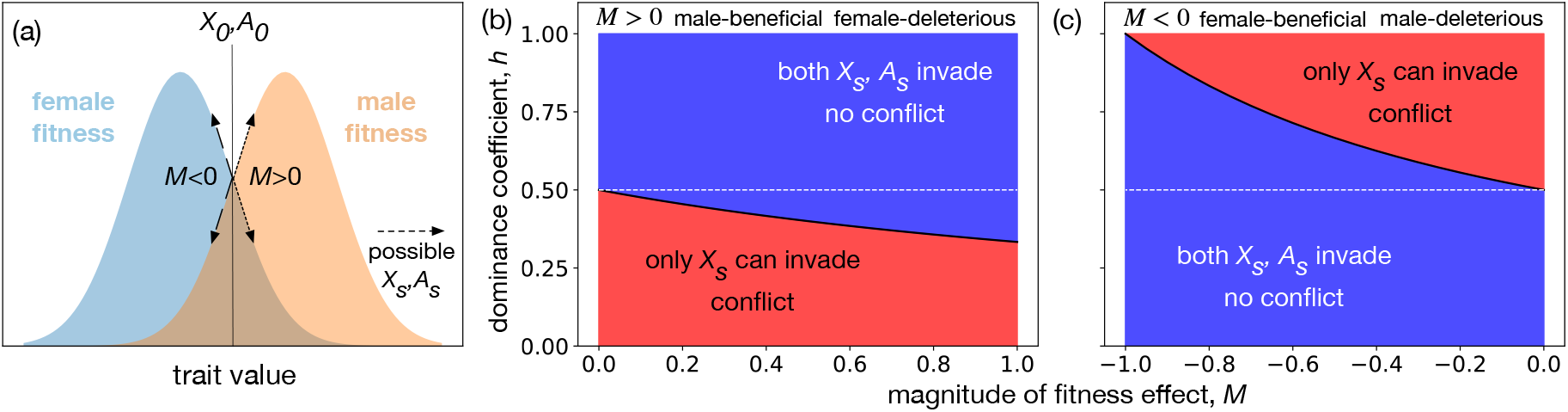
X-autosome conflict over the expression of a sexually antagonistic behaviour. *(a)* We visualise a trait that has single, separate optima in males and females. Starting from a scenario where the resident alleles (*X*_0_, *A*_0_) have identical fitnesses in both sexes (vertical line in the middle), we model the invasion of a mutation which pushes the trait either closer to the female optimum (thereby pushing it farther away from the male optimum, left), or vice versa. Panel inspired by Fig 1a, (Schenkel et al., 2018). *(b)* When an allele is male-beneficial (*M >* 0, equiv. female-deleterious), its dominance has to be low enough to produce a situation where an allele with these effects would go extinct on an autosome, but spread on an X-chromosome. The upper bound on dominance is always lower than 0.5, making it necessarily recessive, and it becomes lower as the magnitude of fitness effects *M* increases (see Ineq. 25 and the sentence after it). *(c)* When the allele is instead female-beneficial (*M <* 0, equiv. male-deleterious), its dominance has to be high enough to produce a situation where an allele with these effects would go extinct on an autosome, but spread on an X-chromosome. The lower bound is always larger than 0.5, making it necessarily dominant, and becomes higher as the magnitude of fitness effects *M* increases (see Ineq. 26 and the sentence after it). There is never a situation where a sexually antagonistic behaviour would spread on an autosome, but not on an X-chromosome. **Alt text**: Three panels. (a) depicts two overlapping normal distributions for male and female fitness as a function of trait value. One type of mutation is labelled *M >* 0, and another *M <* 0. Panels (b) and (c) are plots of the dominance coefficient *h* against the magnitude of fitness effect *M*. Both panels show two regions separated by a line. In (b) the line never goes above *h* = 0.5, region above is labelled “both *X*_*s*_, *A*_*s*_ invade, no conflict”, region below is labelled “only *X*_*s*_ can invade, conflict”. In (c) the line never goes below *h* = 0.5, and the labels are flipped, so that the region above the line is labelled “only *X*_*s*_ can invade, conflict”.

While the treatment so far proves that there can be conflict, we have not focused as much (as previous sections) on whether the selfish alleles invade and then go to fixation, or remain at a polymorphic equilibrium with other alleles at the same locus. Indeed, when a polymorphic equilibrium is possible, it is different across the sexes since the fitness effects are different, and also changes when the allele is autosomal vs. X-linked. Rice (1984) computes equilibrium frequencies for all the cases we consider (see his Eqs. 4,5 for autosomal allele equilibria, and Eqs. 7,8 for that of an X-linked allele).

## 4.4 Adopting the framework to particular biological problems

While we advocate for our approach as a general starting point, the impetus to start modelling most likely comes from wanting to test how some novel variable affects the dynamics of conflict-producing alleles. Our framework is designed to be extended and the obvious question is how. In this section, we highlight existing models for a variety of well-studied internal evolutionary conflicts, which may contain the clues for how to proceed.

### 4.4.1 Segregation asymmetry

Relatively straightforward extensions of the segregation distortion model (see SI section S7) show how driving alleles can spread when their transmission distortion effect is limited to only one sex, or further still, when females mate multiply and a sperm-competition cost is explicitly coupled to male-gamete killing. Though we neglect demographic factors in our framework, driving alleles may often be strongly affected by demographic processes as well as the stochasticity that comes with finite population sizes. For instance, Girardin and Débarre (2021) have demonstrated that when driving alleles come with a cost to the organism, it is more difficult for a driving allele to invade or fixate relative to non-demographic models due to local reductions in carrying capacities induced by the organismal cost. Similarly, Paril and Phillips (2022) have shown that driving elements that are predicted to be fittest in infinite population models need not remain the fittest in more demographically explicit models that take stochasticity and spatial structure into consideration (also see Kläy et al. (2026) for another example of effects particular to stochastic models). Once a meiotic driver has established itself, alleles at unlinked loci that only experience the fitness cost (and not any benefit) are incentivised to suppress the driver – (Scott and West, 2019) provide a broad treatment of this “arms race” between selfish alleles and their suppressors.

Sometimes, however, the question of interest is whether a segregation distorter can invade a population that already contains a driving allele. For example, variants of the autosomal gamete-killing *t*-haplotype that carry recessive lethals tend to be more frequent than those that produce recessive sterility (Klein et al., 1984). One explanation is that inviable embryos cost a mother less resources to produce than what sterile but fully viable progeny might, with those saved resources reallocated to future reproductive output. Charlesworth (1994) and Munasinghe and Brandvain (2024) demonstrate how multiple selfish alleles can be incorporated into a population genetic framework.

While the meiotic drive-type systems above are usually characterised by single multi-genic haplotypes, other methods are necessary to deal with the cross-genome behaviour of transposons. Early transposon models (Hickey, 1982) are closely related to ours, but much theoretical attention has also been paid to finding the equilibrium copy number distribution of transposons in a genome, where a balance can be struck between their transposition activity, and the negative fitness consequences of having too many of them. This was first studied via the diffusion approximation (Charlesworth and Charlesworth, 1983), another core method of population genetics (that we do not use here, but see e.g. Ch. 4-5 of Ewens (1969) or Ch. 15 of Otto and Day (2011)). Such models have yielded insights on their distribution (Ohta, 1984; Sawyer and Hartl, 1986), interaction with mode of reproduction (Dolgin and Charlesworth, 2006), and interactions between different transposon families within the same genome (Brookfield, 1996; Le Rouzic et al., 2007; Xue and Goldenfeld, 2016). In more recent work, Omole and Czuppon (2025) have used this well-developed theory to explain some curious statistical aspects of the copy number distribution.

### 4.4.2 Relatedness asymmetry

Our model of genomic imprinting is merely a caricature designed to illustrate the cause of the conflict. However, more biologically realistic models exist. We assumed that all fitness-affecting interactions occur between individuals that share a maternally inherited allele. This assumption can be relaxed by allowing the interaction structure between siblings to emerge from a population’s mating system. Considering mammalian pregnancy and interactions between siblings in the womb, monogamy, where all offspring in the womb have the same father, can be compared with polyandry, where offspring potentially now have unrelated fathers (Spencer et al., 1998). We also assumed that mating was random, such that each individual was equally likely to mate with another individual. Populations structured across patches or islands violate this assumption. Fortunately, population genetic theory is equipped to deal with this problem, using frameworks such as the “infinite island model”. Van Cleve et al. (2010) show how such complexities can be addressed in the context of the evolution of imprinting. Assortment, i.e. relatedness, asymmetries between paternally- and maternally-inherited parts of the genome have also been long-expected to affect haplodiploids, in particular social insects (Queller, 2003). Dobata and Tsuji (2012) have constructed a quantitative genetics-based model of one such problem, showing that genomic imprinting can evolve on any gene affecting whether or not an immature female develops into a queen.

Substantially more complicated extensions using population genetics are harder to come by, since working with the relevant expressions quickly become cumbersome (see e.g. Gardner et al., 2007). This has prompted the usage of other methods more well-suited to such problems; we provide pointers to these methods in the next subsection “Other modelling approaches and literatures”.

### 4.4.3 Class-linkage asymmetry

This kind of conflict is well-represented in the literature, with the class-structure under consideration usually being that of males and females. While the example mathematical model we built can be applied equally well to the Z-chromosome of a ZZ/ZW sex-determination system, a different perspective is offered by cytoplasmic genetic elements. This is because, in most taxa, they do not pass through both males and females like an X (or Z), but are transmitted purely by females. The consequence of this class-linkage contrast with the autosomes is illustrated in extreme fashion by the arthropod endosymbiont *Wolbachia*, which employs a myriad of strategies to manipulate host reproduction (Werren, 1997; Brenninger et al., 2025). A subset of these strategies involve sex-ratio manipulation, placing them in conflict with organismal interests. Let us focus on one strategy known as “male-killing”: infected male offspring that would otherwise develop normally, die during early life stages (Hurst et al., 1999). This is beneficial from the endosymbiont perspective because females are released from sibling competition with brothers (who are likely to carry the same *Wolbachia* strain), which are evolutionary dead-ends for *Wolbachia* genes (Brenninger et al., 2025). However, this is potentially costly from the perspective of a host autosome, since it distorts the population and primary sex ratio, which are under well-understood selection pressures (see e.g. the book-length treatment by West (2009)). This, as well as other sex-ratio distortion strategies, have been modelled by various authors, showing that this conflict can lead to compensatory sex ratio shifts by the autosomes, polymorphisms in primary sex ratio, population-level outcomes like extinction, etc. (Werren, 1987; Taylor, 1990a; Hatcher and Dunn, 1995).

Prioritisation of female transmission at the expense of male allocation due to class-linkage is a strategy also observed in mitochondria (e.g. the widespread induction of male sterility in hermaphroditic plants by mtDNA; Frank, 1989). Given their shared transmission dynamics, the logic underlying the aforementioned *Wolbachia* models can largely be ported over to models of mitochondria. Clear theoretical treatments of mito-nuclear conflict over male function are presented by Dapper et al. (2023) and Connallon et al. (2018).

A sex-chromosome analogue of maternally-transmitted endosymbionts is provided by the Y- or W-chromosomes, which are exclusively transmitted by (but unlike the endosymbionts also only to) one sex. Nevertheless, the asymmetry in class-linkage can lead to conflict between the Y/W and autosomes, and the Y-chromosome has been shown to be (i) an area of rapid evolution of genes with sex-specific effect (Kaufmann et al., 2021), and (ii) enriched for genes that are male-beneficial and possibly involved in sex-chromosome meiotic drive (reviewed in Bachtrog (2020)).

Lastly, Muralidhar (2019) studies the effect sexual selection may have in facilitating X-autosome conflict, by focusing on the evolution of selfish mating preferences coded for by alleles on sex chromosomes (an X in particular, or Z). If a trait is beneficial to females but deleterious in males (sexually antagonistic), X-linked preferences can evolve which cause females to choose these trait-expressing males – even if they are low-quality – only because the same trait-causing alleles lead to high quality for their female offspring. The female-linkage of an X chromosome thus allows the invasion of selfish mating preferences that would not be able to invade if coded for by autosomal alleles.

## Discussion

### 5.1 Similarities and differences between the three models

The broad structure of all of our models is similar. The assumptions in each model determined organismal interests by displaying when an allele can be termed deleterious (non-driving, non-imprinting, or autosomal versions of alleles with *s >* 0, *b <* 2*c*, or *M* (1 − *γ*) *<* 0 respectively always went extinct). Each set of assumptions included an asymmetry: haploids can compete within diploids but never the other way around, maternal siblings interact and not paternal, and X-chromosomes are inherited differently than autosomes. An organismally deleterious element that would otherwise have been purged by selection is able to persist by exploiting this asymmetry: transmitting itself more often than another allele in heterozygotes (*k >* 1*/*2), silencing itself when paternally inherited (*m >* 0), and being X-linked and optimising its effect on females (*h >* 1*/*2 when female-beneficial, and *h <* 1*/*2 when female-deleterious). Across all cases, it is these latter elements that are in conflict with the organism, and these that we term selfish.

There are also important differences between the models. First, the kinds of life-cycles where each type of conflict can arise are not the same – transmission asymmetries require only diploidy, relatedness asymmetries only require that fitness be affected by social interactions, and class linkage asymmetries only require class-structure and class-linked genetic elements like sex chromosomes.

Second, these conflicts are not equally visible: establishing the presence of a conflict requires one to show that (i) organismal interests oppose the putatively selfish element, and (ii) the selfish element persists by exploiting one of the above asymmetries. Some form of comparison of fitnesses is hence necessary, but not always easily made. For conflicts due to segregation asymmetry, both (i) and (ii) are *relatively* straightforward once a potential selfish allele has been identified, since one need only show that heterozygotes carrying the element have lower organismal-level reproductive output than the wild type (this implies that a non-driving version, for instance, would be purged from the population in the absence of any other counteracting effects). However, for an imprinting allele, one would have to find (or construct) an un-imprinted allele with the same fitness effects and then evaluate whether it goes extinct when all other aspects of the problem (like the form of assortment) are held constant. Such ‘counterfactual’ genetic constructs become relevant also under class-linkage conflicts; a direct recipe to demonstrate conflict would be to alter the proportion of generations an allele spends in a particular class (e.g. manipulate an autosomal locus to segregate like a Y chromosome), then observe whether allele frequencies or the traits they encode evolve in the expected direction (e.g. Rice, 1998; Prasad et al., 2007; Grieshop et al., 2025).

Our example scenarios were idealized in that they exhibited only one of the three types of internal evolutionary conflict. Natural life-cycles, however, are often complex enough to contain the requirements for all three types of conflict, so it is no surprise that many selfish genetic elements arise as combinations of more than one type. For example, Naumova et al. (2001)’s analysis of human genome data suggests that some imprinted regions of the genome are also affected by non-Mendelian segregation, producing conflict due to both segregation and relatedness asymmetry. Úbeda and Haig (2004) have since modelled this, showing how population genetics can be used to understand the interaction between sex-specific segregation of an allele (meiotic drive) and sex-specific expression of the trait caused by that allele (imprinting). Mackintosh et al. (2021) model the dynamics of Xchromosome meiotic drive, which causes conflict both due to transmission distortion and also due to class-linkage asymmetry – the former because the driving X-chromosome causes low organismal fitness, and the latter because a driving X produces a female-biased sex ratio (which is selected to be even at autosomal loci; Cobb, 1914; Gardner, 2023). Lastly, Brandvain (2010) models genomic imprinting on both an autosome (as we did) and an X-chromosome, showing that the invasion conditions for an autosomal vs. X-linked imprinted allele need not be identical (compare Eqs. 9a to 13, however see also their Discussion, p. 519 left). This scenario can lead to relatedness-asymmetry conflict like any imprinted allele, but also to class-linkage conflict between the X and autosomes. There are many more examples that we do not cite here, but see Werren et al. (1988b), Burt and Trivers (2006), Gardner and Úbeda (2017), Scott and West (2019), Fromhage and Jennions (2019), and Patten et al. (2023) for work that considers a wide array of conflicts.

### 5.2 Other modelling strategies and literatures

While the population genetics-based models that we have so far promoted are indeed illuminating and valuable, there are at least two reasons why one must be aware of other modelling traditions. First, some biological scenarios might be too complicated for a population genetic model to be analytically (or conceptually) tractable and other methods become necessary. Second, some scientific problems that are closely related to the notion of internal evolutionary conflict have been studied substantially – just with more applied goals in mind. In this section, we give pointers to the relevant bodies of knowledge and how they have been used.

#### 5.2.1 Kin-selection approaches

In this paper, we provide a framework rooted in population genetics theory and derive invasion conditions that depend on genetic parameters like dominance. Evolutionary explanations for the different conflicts have relied sometimes on genetically explicit intuition or selection acting hierarchically, and other times on patterns of social interactions; we chose to model both these scenarios with population genetics. However, inclusive fitness considerations (as introduced by Hamilton, 1964b) lead others to construct kin-selection models (often described as the Taylor-Frank approach; Taylor and Frank (1996)), where the objective is to understand whether a given phenotype is adaptive in light of its effects on others carrying copies of the phenotype-causing allele. The two approaches are largely interchangeable and complement one another, a fact that has become generally accepted after years of intense debate (the literature on this is extensive, for reviews and perspectives see e.g Lion et al. (2011), Kramer and Meunier (2016), and Lehtonen (2016), for a clear introduction see Taylor (1996), and for mathematical treatments see Lehmann et al. (2016) and Ch. 7 of Rousset (2004)). Nevertheless, a given scenario might lend itself more easily to one approach over the other, so the well-read theoretician should equip themself with both these tools.

Gardner and Úbeda (2017) provided a comprehensive classification of – to borrow the terminology they use – intragenomic conflicts based on the divergence of inclusive fitness interests between different genomic regions. Their work serves as an excellent introduction to inclusive fitness arguments, and illustrates how the phenomena we are interested in can be modelled also using kin selection approaches. In particular, their work contains models of all the same problems we tackle, only constructed with kin selection benefits in mind: see pp. 4-5 of their Supplementary Information for a model of genomic imprinting (they refer to it as an example of “origin conflict”), pp. 5-7 of the SI for a meiotic drive model (referred to as an example of “destination conflict”), and Box 4 for a model of X-autosome conflict over the expression of a sexually antagonistic phenotype (referred to as an example of “situation conflict”).

In fact, Gardner and Úbeda’s (2017) classification (origin, destination, situation) is in agreement with ours. Origin and destination conflict arise due to relatedness asymmetries: origin when a relatedness asymmetry occurs with respect to non-descendant kin, destination conflict when it occurs with respect to descendant kin. We group both of these into one category of any conflict that can be caused due to any relatedness asymmetry, except at the stage of gamete production. Departures from Mendelian segregation at one locus (meiotic drive), and across many loci (transposons) receive their own category, since we find that it is a more natural grouping of phenomena. Situation conflict occurs when two genes have different reproductive values across two classes, and we retain this class. Note, however, that they do not speak in terms of reproductive value – they instead say that the conflicting genes disagree about “context”; the effect of this context must be formalised, as they do in their Box 4, with class reproductive values.

#### 5.2.2 Synthetic gene drives

The study of segregation distortion, and therefore conflict due to segregation asymmetry, has in particular received significant attention as a result of the idea of using such internal evolutionary conflicts to artificially manipulate populations (e.g. mosquitoes carrying Malaria, or invasive species). Originating almost eight decades ago (Vanderplank, 1947; Curtis, 1968), this idea has now given rise to an active area of research (reviewed in parts by e.g. Dhole et al. (2020) and Rode et al. (2019)). Pushing far beyond the simple models we have studied in the present manuscript, synthetic gene drive research has considered effects such as underdominance (Davis et al., 2001; Magori and Gould, 2006), spatial structure (Tanaka et al., 2017; Girardin and Débarre, 2021), inbreeding (Beaghton and Burt, 2022), and many other processes on the success of the driving allele (theoretical aspects reviewed by de Jong (2017), for a recent mini-review see the Introduction of Kläy et al. (2025)). For example, consider the insight that underdominance of fitness effects at a single locus leads to the existence of an invasion barrier for the spread of the lower-fitness allele (also known as “threshold-dependent drives”; reviewed by Leftwich et al. (2018)), such that it can spread only when it begins at a high enough initial frequency. This is a favourable property for a human-introduced driving allele, since it has been shown that gene drives affected by such invasion barriers are spatially self-limiting, *i.e*. they cannot take over a neighbouring population unless migration is so strong that it overcomes the invasion barrier (Dhole et al., 2018). This result could have interesting consequences for naturally occuring systems, e.g. range-restricted *D. melanogaster* segregation distorters might be associated to underdominance. Numerous such insights, obtained with a focus on translational research, can shed further light on the fundamental biology of segregation distorters.

## Conclusions

There are many reasons to be interested in the study of internal evolutionary conflicts (Ågren and Patten, 2025). The logic is “intellectually delicious” and their existence “challenge just about every presumption you might have had about evolution and natural selection” ((Hurst, 2025) p. 189). At a conceptual level, their very existence produces fractures in the image of an organism as a fitness-maximising entity (Patten et al., 2023). Some are particularly important as they have become major drivers of evolution across taxa: transposable elements make up as much as 80% of the host genome in some plant species (Wegrzyn et al., 2013), and have been postulated to be major sources of mutation and variation for the host populations that they inhabit (Bourque et al., 2018). Others are of medical relevance, including genomic imprinting which may provide an evolutionary explanation for some human disorders (Úbeda, 2008).

In this paper, we have provided an introduction to the theory of internal evolutionary conflicts by means of three example mathematical models. These models serve not only as tools to make our arguments precise, but also to introduce the reader to the process of building and analysing a mathematical model. While the techniques we use here are well-known, they nonetheless provide fundamental insights. We also offer a definition of an internal evolutionary conflict, and use it to contrast many biological phenomena that are invoked in this context. Our discussion of other modelling approaches then allows one to gain further modelling expertise if they wish to do so.

An integral part of continuing this work will be to maintain close communication between theoretical and empirical advances. Indeed, the persistence of debates over definitions and the time it has taken to prove general claims in this field is a testament to how diverse such phenomena can be in their biological details. While this presentation does not claim to be exhaustive, we are hopeful that it will accelerate further understanding.

## Data Availability Statement

All results are produced by analysis of mathematical models. Scripts used to produce figures and perform the numerical simulations in the supplementary can be found at the following GitHub repository: https://github.com/gauravathreya/how-to-conflicts.git

## Acknowledgements

The authors thank the organising committee of the ESEB Special Topic Network on Internal Conflicts and Organismal Adaptation for organising the October 2024 workshop in Groningen where this work began. We also thank the Kokkonuts for detailed comments and valuable discussion.

## Supplementary Information

### Outline of the Supplementary

In this Supplementary Information document, we provide additional context and support for arguments or claims that we make in the main text. Due to the high diversity of models that exist of transmission distortion, we show in Section S7 how our simplistic model can be connected to more involved models that incorporate specific aspects of empirically important, well-studied examples of transmission distortion. In Section S8, we show that the Price Equation (Price, 1972) can be, in principle, used in similar ways and to similar ends as the population genetics models. Section S9 discusses how one can produce invasion criteria when an allele has different fitness effects across two classes (here, males and females). Finally, in Section S10 we demonstrate that class reproductive values affect invasion criteria in a class-structured population, showing that conflict due to class linkage does, in fact, arise due to class linkage.

### S7 Some extensions

In many organisms, the expression and strength of drive is sex-specific (Courret et al., 2023). In this section, we demonstrate how such sex-specific drive can be modelled analytically in a population genetic framework. Just as in the main text, we formulate simple one-locus, two-allele models and compute the equilibrium frequency of a driving allele that increases its own transmission efficiency but bears an organismal cost. To make the cost to the organism particularly strong, we will assume throughout this section that the driving allele is homozygous lethal, such that amongst adults it can only be found in heterozygotes.

#### S7.1 The mouse *t*-haplotype: homozygous lethality and sex-specific drive expression

The *t*-haplotype complex is a region of an autosomal chromosome in wild mice that is known to have male-limited transmission distortion (Lyon, 2003). Males that are heterozygous for the selfish *t* transmit it to nearly all of their sperm, whereas segregation is fair in females. Additionally, the *t*-haplotype is non-recombining and most variants are homozygous lethal, thus only occurring in heterozygotes (Klein et al., 1984). Here, we extend the main text transmission distortion model to calculate the expected equilibrium frequency of a transmission distorter with *t*-like properties.

Consider an autosomal transmission distorter *E*_*s*_ that is homozygous lethal but is associated with no survival costs in heterozygotes. Suppose male heterozygotes transmit the *E*_*s*_ allele in a fraction *k*_♂_ ∈ (0, 1] of gametes, whereas females transmit the *E*_*s*_ allele in a fraction *k*_♀_ ∈ (0, 1] of gametes. Thus, transmission distortion exists when (*k*_♂_, *k*_♀_)≠ (1*/*2, 1*/*2) and is sex-specific when *k*_♂_≠ *k*_♀_. For example, an element that drives only in males would have *k*_♀_ = 1*/*2, *k*_♂_ *>* 1*/*2. Unlike the models in the main text, we will track the change in *p* here by counting all possible matings that can happen. Assuming random mating, the set of all possible matings and the kinds of offspring they produce can be listed concisely via a mating table (table S1). The total number of genotypes involving *E*_*s*_ alleles in the next generation is calculated by summing up the contributions from each Supplement to Athreya et al., relevant mating combination:

**Table S1.**
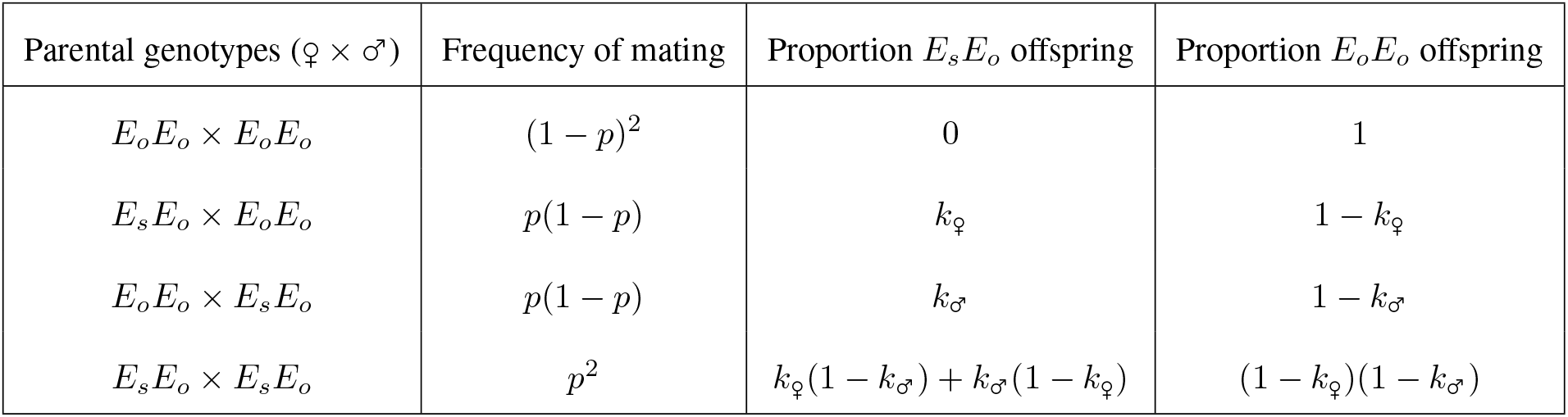
Table of all possible matings and their outcomes for sex-specific drive. *k*_♂_ and *k*_♀_ are sex-specific segregation ratios. *k*_*◦*_ = 0.5 indicates fair (Mendelian) segregation.

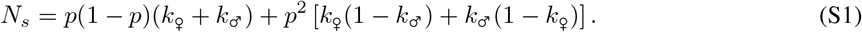

Similarly, the total number of genotypes involving *E*_*o*_ alleles in the next generation is

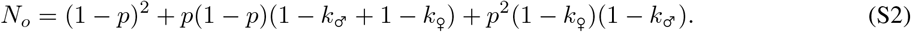

Since *E*_*s*_ is only present in heterozygotes, the total *allele* frequency of *E*_*s*_ in the next generation is thus

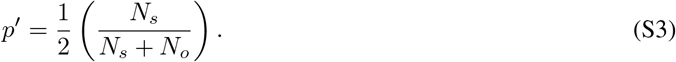

Substituting Eq. S1 and Eq. S2 into Eq. S3 and setting *p*^*′*^ = *p* to find the equilibrium frequency yields, after some lines of algebra, the equilibrium frequencies *p*^∗^ = 0 and

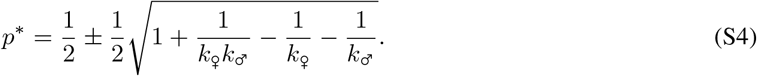

Setting *k*_♀_ = 1*/*2, *k*_♂_ = *m* recovers the predictions of a classic model for the evolution of the *t*-haplotype in mice (Bruck, 1957, Eq. 5)

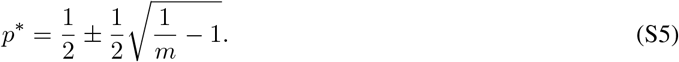

Furthermore, making drive strength the same in both sexes by setting *k*_♀_ = *k*_♂_ = *k* yields the equilibrium frequency

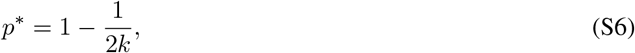

which describes the same outcome as Eq. 8 in the main text and Eq. S23, the same result derived using the Price equation. The numerical value of the equilibrium above differs to main text Eq. 8 and Eq. S23 by a factor of 2*k* because the latter models census organisms at the zygote stage, but Eq. S6 censuses organisms at the adult stage (explained further just before Eq. (S23)).

With sex-specific drive, the deleterious effects of the allele (loss of some offspring due to homozygote lethality) can be offset by driving in just one of the two sexes. In fact, even stronger outcomes are possible — Eq. S4 predicts that a homozygous lethal genetic element could persist even if it drags in one sex as long as it drives sufficiently strongly in the other (*k*_♂_ *<* 0.5, *k*_♀_ *>* 0.5 or vice versa).

At *k*_♀_ = 1*/*2, *k*_♂_ = 0.9, the typical drive strengths for many male-specific driving elements including the *t*-haplotype, Eq. S4 predicts an equilibrium frequency of 0.33 (Fig S1), or 33%. However, empirical measures of the frequency of driving alleles in natural populations consistently find that these alleles are segregating at significantly lower frequencies. For instance, Brand et al., 2015 find that *Segregation Distorter*, a sex-limited autosomal meiotic driver in *Drosophila melanogaster*, is present at a frequency of about 1-8% in natural populations. What may explain such low prevalences? Various ecological factors have been found to affect the predicted equilibrium frequency of meiotic drivers (Lindholm et al., 2016). In the next section, we will see how one such ecological factor, sexual selection via sperm competition, can be incorporated into the population genetic model we have explored in this section.

**Figure S1.**
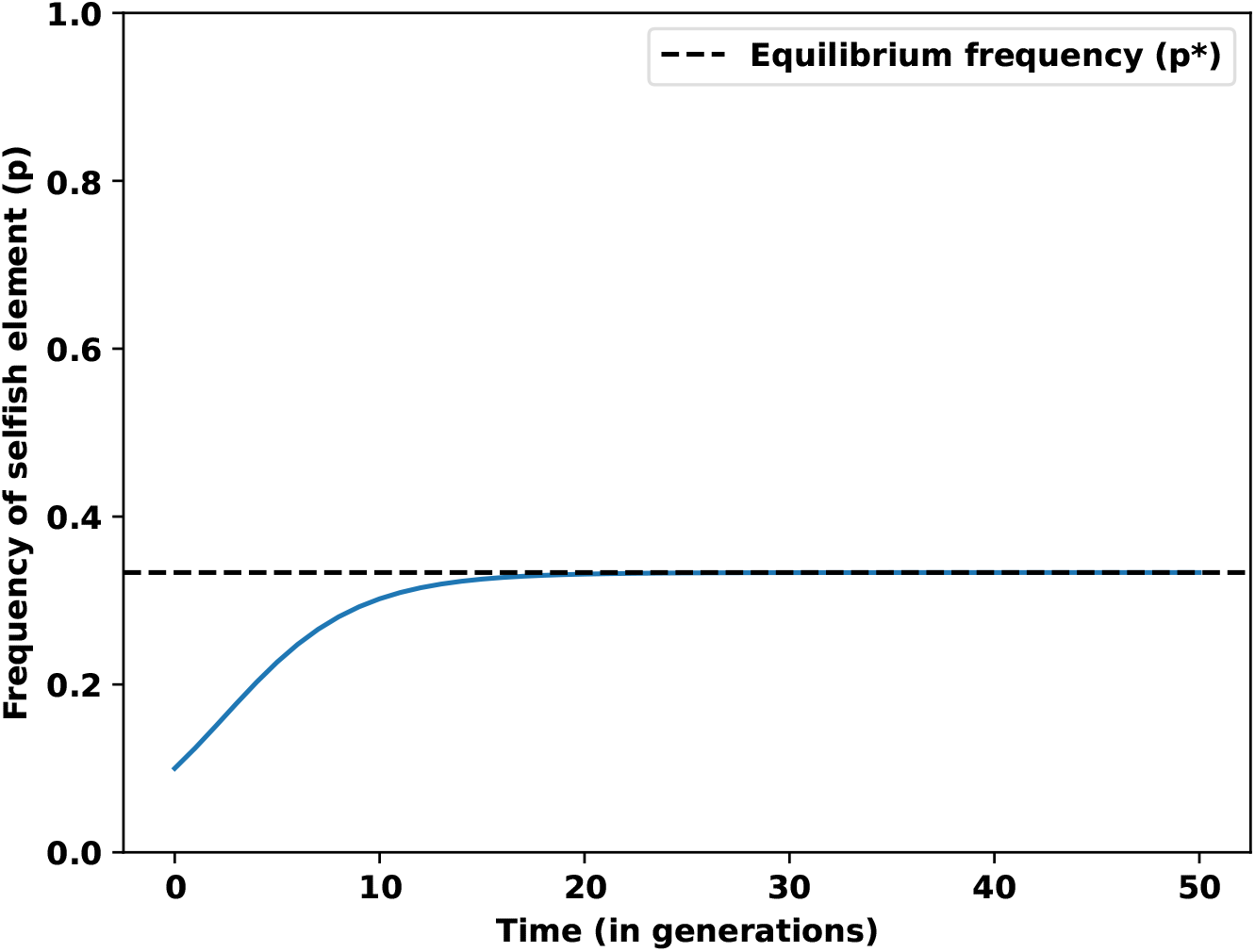
The trajectory of allele frequency over time predicted by our model of sex-specific drive for *t*-haplotype style parameters (*k*_♀_ = 0.5, *k*_♂_ = 0.9, homozygote lethality, and no organismal cost to heterozygotes). Blue line is the recursion Eq. S3, and the dotted line is the equilibrium frequency predicted by Eq. S4. For this set of parameters, *p*^∗^ = 0.33.

#### S7.2 The *Drosophila melanogaster* Segregation Distorter: multiple mating and sperm competition

*Segregation Distorter* (SD) is a selfish gene-complex found on chromosome 2 of *Drosophila melanogaster. SD* variants distort transmission during spermatogenesis, where by some unknown mechanism, they sabotage the development of spermatids that do not carry the *SD* element. The result is that males heterozygous for *SD* produce primarily *SD* bearing sperm, in quantities that often exceed 90% (Larracuente and Presgraves, 2012). While the driving effect is limited to males (see table 5 in Brand et al., 2015 for an estimate of sex-specific drive strengths), irrespective of sex, it is extremely deleterious when present in two copies (i.e. when the organism is homozygous for the element). In many cases, the element is homozygous lethal (Temin and Marthas, 1984; Wong and Holman, 2019).

The model presented in section S7.1 predicts that an *SD* element that is present in 90% of a heterozygote male’s viable sperm (*k*_♂_ = 0.9, *k*_♀_ = 0.5) should be present at a frequency of 0.33 at equilibrium (Fig S1). However, in natural populations, *SD* is typically present at much lower frequencies of about 1-8% (Brand et al., 2015). In a recent study, Keaney et al., 2021 investigated the role sexual selection could play in reducing the frequency of *SD* elements. In the case of *SD-5*, a North American variant of the complex, Keaney et al. (2021) experimentally found that *D. melanogaster* females are more likely to remate when mated with a male that carried *SD-5*. Furthermore, when females mated multiply, males carrying *SD-5* sired fewer offspring than wildtype males, suggesting that bearing *SD-5* is also associated with costs during sperm competition. To link these results with theory, Keaney et al., 2021 built evolutionary simulations to predict the equilibrium frequency of *SD* in polyandrous populations. Here, we show that one needn’t turn to simulations so quickly. Instead, the model presented in section S7.1 can be extended to allow females to mate twice.

We use the same strategy as the previous section, except that the mating table is now given by table S2. Here, *β*_*i*_ is the probability that a female remates if its first mate had genotype *i* (*s* means genotype *E*_*s*_*E*_*o*_ and *o* means genotype *E*_*o*_*E*_*o*_), and *γ*_*ij*_ is the proportion of offspring sired by the mate of type *i* when *i* was the first mate and *j* was the second mate. We assume that females mate either once or twice, that mating encounter frequencies are given by genotype frequencies, and that females bear no fecundity costs from carrying the selfish allele. The parameters *γ*_*ij*_ and *β*_*i*_ can be thought of as modelling sperm competition. In particular, if the driving allele *E*_*s*_ is a sperm killer, *E*_*s*_*E*_*o*_ males are expected to produce fewer sperm than *E*_*o*_*E*_*o*_ males. As female refractory-period inducing peptides bind to sperm tails (Peng et al., 2005), females may therefore be more likely to remate if they first mated with a *E*_*s*_*E*_*o*_ male (*β*_*s*_ *> β*_*o*_). Indeed, Keaney et al. (2021) empirically found that *β*_*o*_ ≈ 0.3 and *β*_*s*_ ≈ 0.75 when *D. melanogaster* females were given the opportunity to remate several days after having mated with *SD-5* or non-driving chromosome carrying males, respectively. In the event of multiple mating, the heterozygote may also face a systematic disadvantage under sperm competition, due to having a smaller ejaculate. While *D. melanogaster* exhibit last-male sperm precedence (Morrow et al., 2005) (*i.e*. a female’s most recent mate sires most of her offspring), Keaney et al. (2021) additionally found that carrying *SD-5* reduces this advantage, relative to a wildtype male in the second mating role. That is, for males in the first mating role *γ*_*so*_ *< γ*_*os*_ *<* 0.5.

**Table S2.**
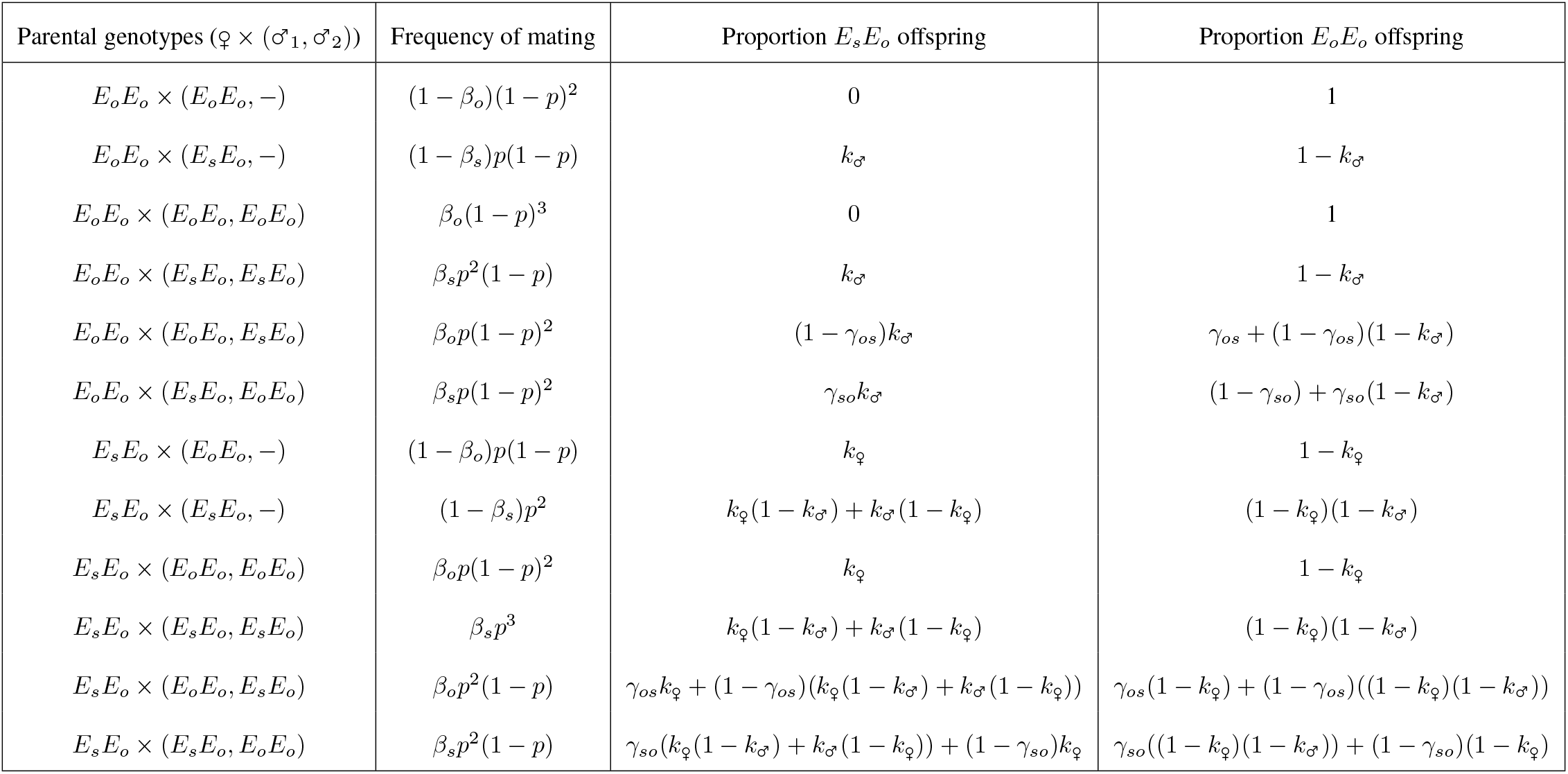
Table of all possible matings and their outssscomes when a female is allowed to mate (at most) twice. ♂_*i*_ indicates that the male is the *i*^th^ mate, and − indicates that the female does not remate. *β*_*i*_ is the probability that a female remates if its first mate had genotype *i* (*s* means genotype *E*_*s*_*E*_*o*_ and *o* means genotype *E*_*o*_*E*_*o*_), and *γ*_*ij*_ is the proportion of offspring sired by the mate of type *i* when *i* was the first mate and *j* was the second mate.

As before, we can now calculate the frequency of the selfish allele *E*_*s*_ in the next generation by computing *N*_*s*_ and *N*_*o*_, the total number of *E*_*s*_ and *E*_*o*_ alleles respectively, from the mating table S2, and then substituting the expressions into Eq. S3. We plot the predicted equilibrium frequencies as a function of *γ*_*ij*_ for the parameters corresponding to Keaney et al. (2021) in Fig S2.

**Figure S2.**
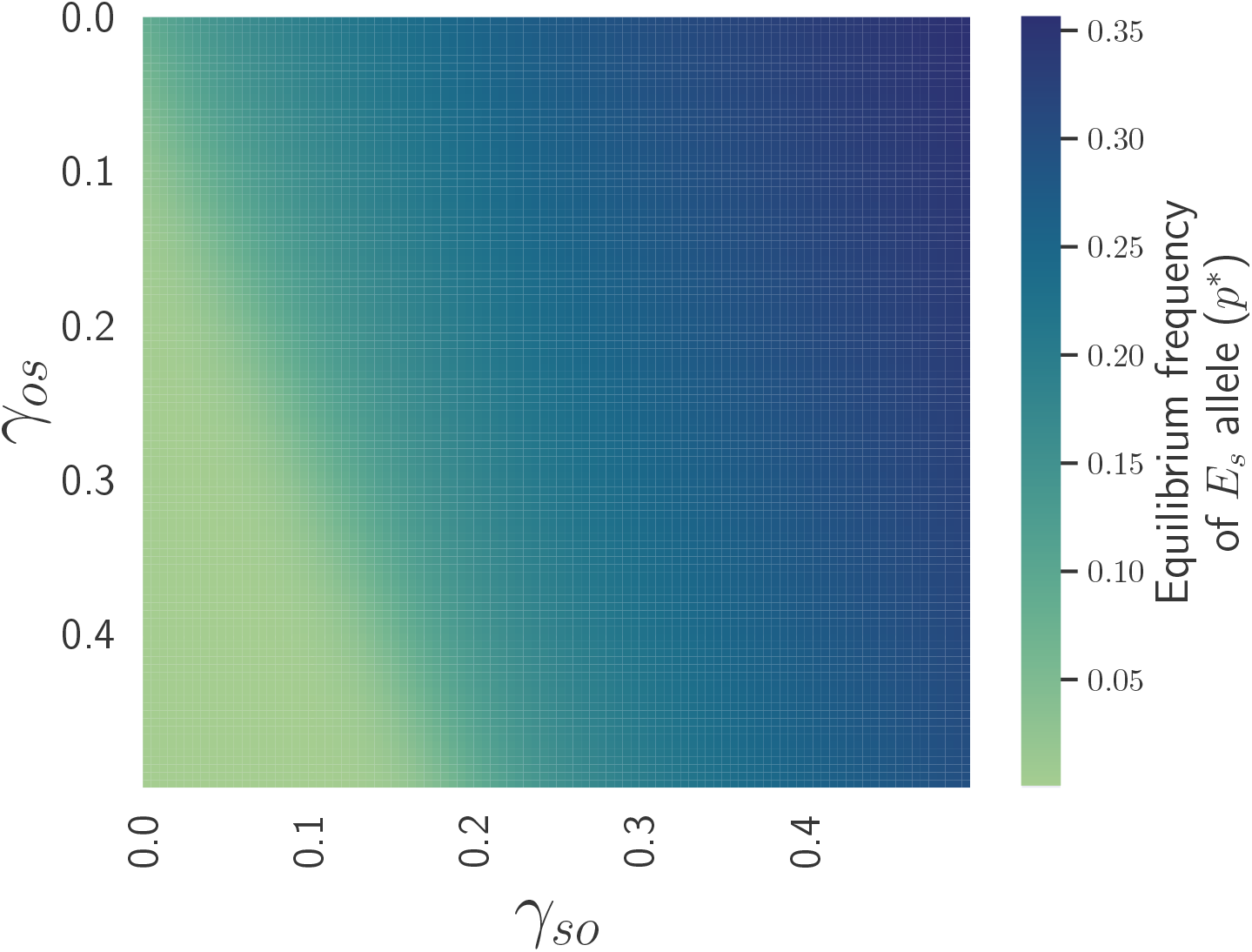
Equilibrium frequencies of the driving allele when a female is allowed to mate twice. *γ*_*SO*_ is the proportion of offspring sired by a SD-carrying male in the first mating role, with a wild-type male in the second mating role. *γ*_*OS*_ is the proportion of offspring sired by a wild-type male in the first mating role, with a SD carrying male in the second mating role. Fixed parameters are adapted from Keaney et al., 2021: we set *β*_*o*_ = 0.3, *β*_*s*_ = 0.75, *k*_♀_ = 0.5, *k*_♂_ = 0.944.

Figure S2 shows that sperm competition, here modelled via *γ*_*ij*_, strongly impacts the predicted equilibrium frequency of the driving allele in our model. Predictably, the equilibrium frequency of the driving allele reduces as the cost of the driving allele to sperm competitive ability increases (*γ*_*so*_ decreases, *γ*_*o*_*s* increases – bottom left of Fig S2). In *D. melanogaster, SD-5* is a known sex-specific meiotic driver with homozygote lethality. Keaney et al. (2021) (in their supplementary) empirically estimate *γ*_*so*_ = 0.01, *γ*_*os*_ = 0.08, *β*_*o*_ = 0.3, *β*_*s*_ = 0.75, *k*_♀_ = 0.5, *k*_♂_ = 0.944 for this system (see the table “estimated proportion of offspring sired by SD/+ or control males, when they mated first.” in https://tomkeaney.github.io/SD_sexual_selection/Empirical_analysis.html#Measuring_P1 for *γ*_*ij*_ values). Our model predicts that the equilibrium frequency of the *SD-5* allele should then be *p*^∗^ = 0.025, or 2.5%. This estimate is roughly consistent with previous empirical studies suggesting that the global frequency of *SD* chromosomes is between 1-8% (Brand et al., 2015). The equilibrium frequency predicted by the simpler model Eq. S4 is about 0.378 (Bruck, 1957), a much higher value that hasn’t been observed in natural populations.

### S8 Internal Evolutionary Conflicts in the Context of the Price Equation

Our definition of an internal evolutionary conflict requires divergent evolutionary interests between a selfish allele and the larger organism within which it resides. This hierarchical structure is the focus of multilevel selection models, which heavily lean upon Price’s general theorem of selection (Price, 1970; Price, 1972). The theorem, often referred to as the Price equation, provides a general description of the expected evolutionary change in a population (Frank, 2012; Gardner, 2020). Within-individual selection has been shown to be fully compatible with this framework, which has been used for both identifying internal evolutionary conflicts and their evolutionary outcomes (Okasha, 2006; Patten et al., 2023; Patten, 2025). Here we demonstrate why the Price approach is valuable for modelling internal evolutionary conflict, using a transmission distortion example. We model the same problem as presented in the main text, recapitulating the result using the shift in focus that the Price approach demands.

Before we get there, it is important to clarify how trait distortion fits into Price’s theorem. These conflicts most often occur when interactions between relatives affect fitness, and the optimal social behaviour differs between agents within an organism. To effectively capture fitness effects that come about due to interactions between organisms, levels of organisation above that of the organism need to be modelled. Such extensions of the Price equation exist (reviewed in (Lehtonen, 2016)), and have been used to model the evolution of behaviours like spite and selfishness (Hamilton, 1970). A full treatment of the problem is beyond the scope of this manuscript, but is a fruitful extension that we encourage.

Returning to the problem at hand, there are two key parts to Price’s argument. First, consider the relative number of descendants that specific categories in a population produce. Second, consider the fidelity with which members of these categories transmit information to descendants (where information can be alleles, genotypes, trait values etc.).

More formally, let the frequency of members in category *i* across the population be *q*_*i*_ and the average of the value you’re interested in tracking within the category be *z*_*i*_. Selection determines the fraction of total descendants 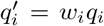 produced by members of category *i*, where *w*_*i*_ is the relative fitness of members of category *i*. The mean value found in these descendants is 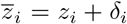, where *δ* is a deviation caused by imperfect transmission. The crucial difference between Price’s logic and classical population genetics is what 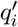 represents. It is not the frequency of descendants of category *i* in the next time-point, but the frequency of individuals at the next time point that descend from individuals currently in category *i*. Putting this all together, an instructive form of Price’s theorem for students of internal evolutionary conflict is

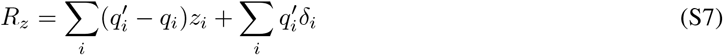

where *R*_*z*_ is the evolutionary change in the average value of *z* across all categories in the population (see (Frank, 1997; Frank, 2012) for more details on this variant of the theorem). The first term is the change due to selection operating between categories. The second (with *δ*_*i*_) is the change caused by imperfect transmission of Supplement to Athreya et al., ancestral values to descendants, and importantly for our purposes, can include selection operating *within* members of categories. It has been suggested that when the signs of the first and second terms in Eq. (S7) are opposite, there is evolutionary conflict between entities at different levels of organisation (discussed in (Patten et al., 2023)).

To demonstrate how this variant of the theorem can be applied, we model the fate of a rare selfish allele *E*_*s*_ that benefits via transmission distortion in a population initially comprised of the wildtype variant *E*_*o*_. Let the form of distortion be segregation distortion and for simplicity, consider only fitness effects at its diploid locus. Let the frequency of the *E*_*s*_ allele be *p*. Following Price’s logic, we first need to specify the categories to track in the population; let’s make these the invading allele *E*_*s*_ and the resident wild type allele *E*_*o*_. Set their values to *z*_*s*_ = 1 and *z*_*o*_ = 0, such that the mean value in the population is 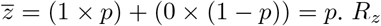 therefore tracks the change in *p*. Since we assume the driving allele is homozygous lethal (as is commonly the case for observable gamete-killer drives), the viability of *E*_*s*_*E*_*s*_ homozygotes is zero. We set the viability of all other genotypes to 1 and assume no effects of the locus on any other organismal-level fitness components. Considering only organismal-level selection, the marginal absolute fitnesses of the *E*_*s*_ and *E*_*o*_ alleles are, respectively

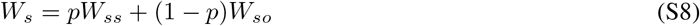

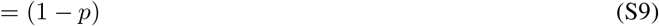

and

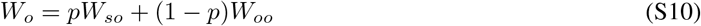

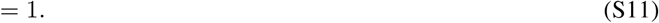

The mean fitness of the two alleles is

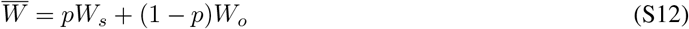

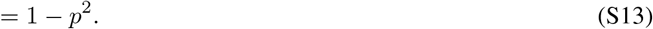

With relative fitness 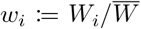, we have all the information required to calculate 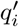, the fraction of the adult population descended from category *i*. Entering the relevant absolute fitnesses lets us calculate 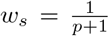 and 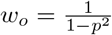 Thus, considering only the first term in Eq. (S7) we find

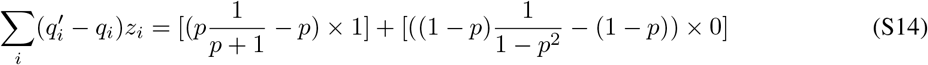

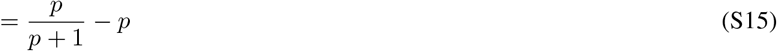

which is negative or zero for all possible frequencies of the *E*_*s*_ allele. That is, selection *between* individuals never favours the driving *E*_*s*_ allele.

We now turn to the change in allele frequency caused by transmission distortion. In the absence of mutation and other non-adaptive effects, *δ*_*i*_ represents a departure of strength *d* from Mendelian segregation caused by transmission distortion. *d* = 0 indicates fair segregation, whereas positive *d* produces drive. Note that *d* is called drive strength in the main text, and is related to the transmission distortion parameter *k* in the main text by *d* = 2*k* − 1. In other words, *k* determines the fraction of alleles transmitted by heterozygous genotypes, whereas *d* the transmission fidelity of an allele across generations. Conveniently, 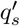 tells us the frequency of *E*_*s*_ alleles that appear alongside *E*_*o*_ alleles in heterozygotes (as *E*_*s*_*E*_*s*_ genotypes have all died), and therefore, the proportion of alleles threatened by transmission distortion. Under random mating amongst the surviving genotypes and drive operating in the heterozygotes of both sexes, the partial change in allele frequencies caused by transmission bias is

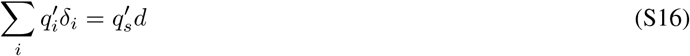

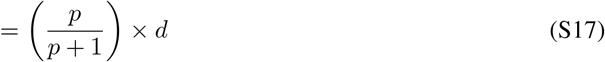

which can be interpreted as the frequency of *E*_*o*_ alleles inhabiting heterozygotes that are converted or replaced in the gamete pool by *E*_*s*_ alleles. When *d >* 0, the sign of *δ*_*i*_ is positive. Finally, the total change in the frequency of the driving allele *E*_*s*_ from one generation to the next is

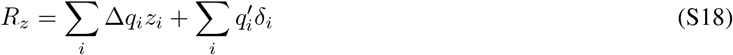

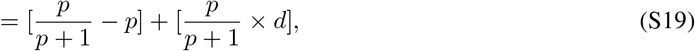

which can be rearranged to find

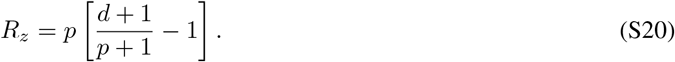

Equilibrium frequencies *p*^⋆^ for the *E*_*s*_ allele are found by setting *R*_*z*_ = 0. This yields that either *p*^∗^ = 0 or

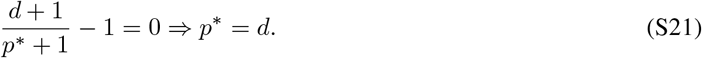

To see that Eq. S21 predicts the same equilibrium frequency as the equation we arrived at in the main text, substitute *s* = *W, h* = 0 into Eq. 8 from the main text. This yields

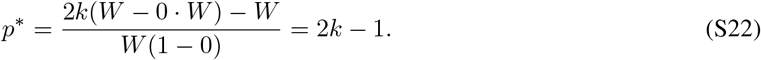

Recalling now that *d* = 2*k* − 1 makes the connection complete. To connect this approach to the sex-specific drive equilibrium frequency Eq. S6, we we must convert the *p*^∗^ of Eq. S21, which tells us the equilibrium frequency in the zygote stage, to the equilibrium predicted amongst adults. To get there, just remove those zygotes that perish Supplement to Athreya et al., before adulthood

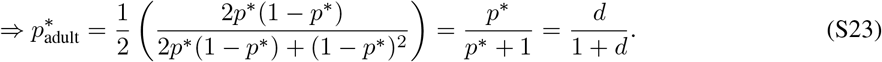

Substituting *d* = 2*k* − 1 now yields Eq. S6 as promised.

### S9 Invasion criteria for systems of 2 recurrence equations

For many applications, one might wish to consider a class-structured population where alleles have different fitness effects across the classes. In the example of X-autosome conflict over sexual antagonism in the main text, it is similarly necessary to construct a system of two recurrence equations, one for the allele frequency in males and another for allele frequency in females. To compute invasion criteria in such a scenario, it is not possible to directly use the methods of Box 2 since they are meant for populations that can be described by one-dimensional equations. However, a small generalisation is sufficient: instead of computing one derivative, we now need to compute a 2 *×* 2 matrix of derivatives called the *Jacobian* matrix.

Consider a set of two recurrence equations describing the allele frequency dynamics of an allele *E*_*s*_ at a locus where most of the population is homozygous for a resident allele *E*_0_. Suppose the dynamics is given by

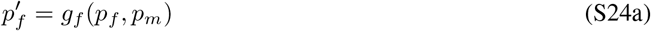

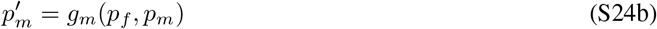

where *p*_*f*_, *p*_*m*_ are the current allele frequency of *E*_*s*_, 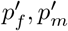 are next-generation frequencies of the same allele, and the functions *g*_*f*_ (*p*_*f*_, *p*_*m*_), *g*_*m*_(*p*_*f*_, *p*_*m*_), in general different to each other, determine how these frequencies change from one timestep to the next. We impose only that, once the rare allele *E*_*s*_ has arisen due to mutation, migration, etc., no more events take place that cause further influx of *E*_*s*_. In other words, if *E*_*s*_ goes extinct, it cannot bounce back – thereby making (*p*_*f*_, *p*_*m*_) = (0, 0) an equilibrium.

To understand when *E*_*s*_ can invade, we need to again check when a small quantity of *E*_*s*_ carrying organisms can increase from rarity. If the allele frequencies *p*_*f*_, *p*_*m*_ are indeed very small, we can assume that terms in the functions *g*_*i*_() that are 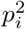 or higher power are negligible, since multiplying a number between 0 and 1 by itself only makes it smaller. Once we have done this, we are left with a set of equations of the form

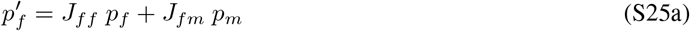

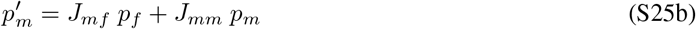

which can be rephrased in the language of matrices as

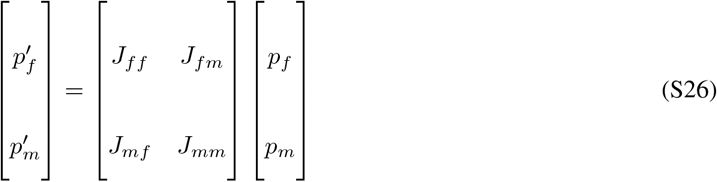

where the matrix made up of the numbers *J*_*ij*_, *i, j* ∈ {*m, f*} is called the Jacobian matrix associated to this system of equations.

We can also arrive at this matrix equation via another route, i.e. to “linearise” Eq. (S24), which amounts to performing a Taylor expansion of Eq.S24 around the equilibrium (*p*_*f*_, *p*_*m*_) = (0, 0), and assume that terms in 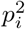 or higher powers of *p*_*i*_ are negligible exactly as we initially did above. The extra knowledge one gains from this is that the numbers *J*_*ij*_ are the first-order partial derivatives of the functions *g*_*f*_ and *g*_*m*_, and are given by

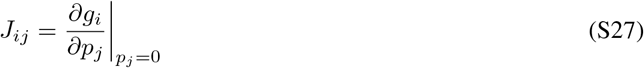

The allele *E*_*s*_ invades when the matrix *J* has a dominant eigenvalue larger than 1. The eigenvalues are defined as the roots *λ* of the polynomial det(*J* − *λI*) = 0, known as the characteristic polynomial associated to *J*. When *J* is a 2×2 matrix, it can be shown that the characteristic polynomial is given by

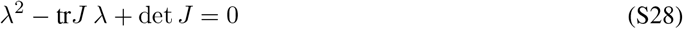

where tr*J* is the *trace* of the matrix *J*, defined as *J*_*ff*_ + *J*_*mm*_, the sum along the diagonal in Eq. (S26); and det *J* := *J*_*ff*_ *J*_*mm*_ − *J*_*fm*_*J*_*mf*_ is called the determinant of *J*, and is also calculable from just the matrix entries. In the simple case of 2 classes and 2×2 matrix, the roots of the characteristic polynomial (S28) can be found using the quadratic formula, and are given by

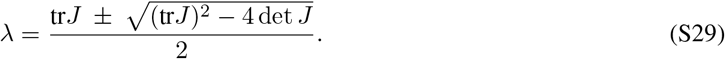

There are two roots, one that can be found by adding the square-root term (the + in the *±*) in the numerator, and another that can be found by instead subtracting it (the − in the *±*). The dominant eigenvalue is defined as the one with larger magnitude (also when they are complex); which root is larger depends on the problem at hand and the entries of the Jacobian.

For the X-autosome example in the main text, the linearised equation for the dynamics of the autosomal allele *A*_*b*_ takes the form

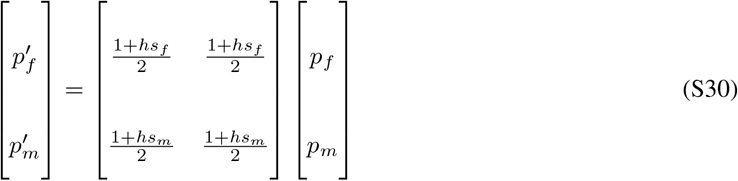

which has the eigenvalues *λ* = 0 and 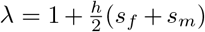, of which the latter is larger, and in addition larger than 1 when *s*_*f*_ + *s*_*m*_ *>* 0.

For the X-linked allele *X*_*b*_, the linearised equations take the form

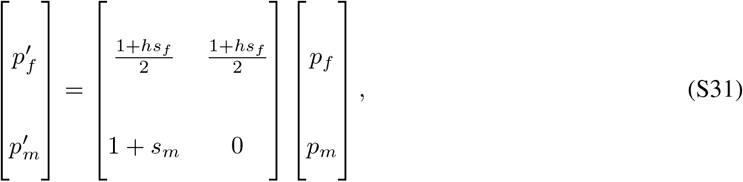

which has eigenvalues given by

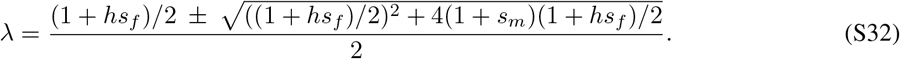

The ‘+’ root is always the larger of the two, and in addition larger than 1, when

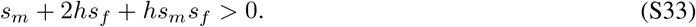

### S10 Class reproductive values affect invasion criteria

In this section we will demonstrate that the invasion of an allele in a class-structured population depends not only on its fitness effects in the different classes, but also on the reproductive values of those classes. We do this by considering a slightly more general version of the above (§S9) model of two sexes, and within this model studying the invasion of an allele with sex-specific effects. The demonstration will proceed in three steps: first we will introduce the notion of class reproductive value and compute it in a simple setting, second we calculate the invasion criterion for an arbitrary allele in a population with separate sexes, and then lastly we will combine the two expressions to prove the claim made above.

#### S10.1 Class reproductive value

Class reproductive values are so named since they indicate how much of the alleles in the future are descended from the alleles found in a specific class at the present time. It becomes important for evolutionary arguments when there is meaningful class-structure in the population of interest, such as separate sexes, ages, ploidy, etc. The classes may differ in their survival rates, fecundities, or more broadly in an allele’s fitness effect on organisms of that class.

To define class reproductive value, we will need to digest a specific matrix first. Consider a locus in organisms of a randomly mating, i.e. panmictic, population with two classes. We will not specify whether this locus is on an autosome, or a sex chromosome, has otherwise equal ploidy in both sexes, maternally transmitted symbiont, etc. Instead, to generalise the idea of “inheritance rules” that can change between loci on differently inherited genetic elements, think of the following square matrix ℐ. ℐ has as many rows and columns as there are classes (2 in the case of females and males), and element ℐ_*ij*_ is the probability that a gamete produced by a class-*i* organism contains an allele inherited from a parent of class *j*.

For example, in the case of alleles going between males and females, ℐ_*ff*_ is the probability that a female-produced gamete contains a female-produced allele (at this locus of interest). In the case of an autosomal allele, the inheritance matrix (call it ℐ_*A*_) is given by

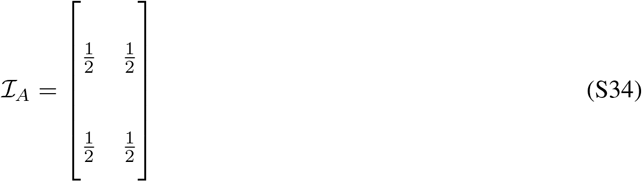

since both males and females are diploid at autosomal loci, any focal male or female received one allele each from its mother and father, and both alleles at a locus are equally likely to make it into a gamete. More intuition is found by thinking about the corresponding matrix for an X-linked locus, which is given by

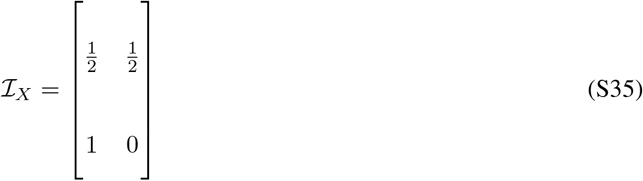

which is 1*/*2 on the top row for the same reason as ℐ_*A*_: females are diploid at an X-linked locus, any focal female received one allele each from its mother and father, and both alleles at a locus are equally likely to make it into a gamete. However the bottom row is different because males are haploid at an X-linked locus, they receive this single allele only from a female and nothing from a male (hence bottom left entry *p*_21_ is 1 and bottom right 0), and as a result they can pass on only their maternally inherited allele. Other inheritance rules can be treated similarly, for example in the case of the maternally transmitted endosymbiont *Wolbachia*,

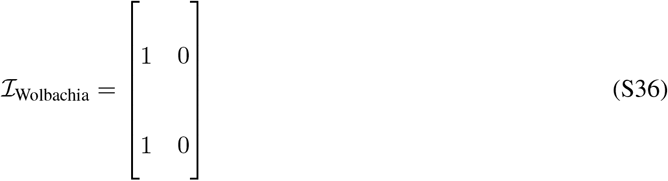

since only females can transmit Wolbachia. As one might have already noticed, the entries along each row of any such matrix ℐ must sum to 1, since the gametes produced by a class *i* organism can only carry an allele that they inherited from some other class – it must have come from *somewhere*. The examples above are of sex-structured populations, where the class structure of consequence is to be male or female; we chose these examples to motivate the matrix ℐ since they are relevant to the X-autosome conflict in the main text. However, the classes can also be ages, or ploidy levels, etc. – one can always speak of the probability ℐ_*ij*_ that, given an allele produced by class-*i*, that it was derived from an organism of class *j*.

To showcase the generality of this section’s claim, we will hereafter use only the fact above that row sums must equal 1, and suppose that the matrix has the following form

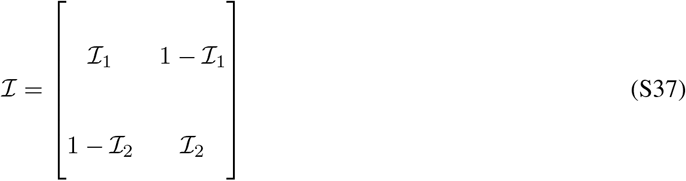

where the number ℐ_*i*_, *i* ∈ {1, 2} denotes the probability that a gamete produced by a class *i* organism contains an allele inherited from a parent of its own class – a kind of autocorrelation of allele transmission among classes. The class reproductive values of each class, *c*_1_ and *c*_2_, are defined as the numbers satisfying the equation

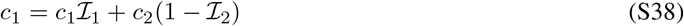

under the constraint that

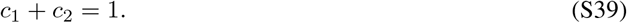

The former equation has an interpretation in a time-recursive sense: alleles produced by class-1 organisms came either from a class 1 parent (with probability ℐ_1_), or from a class 2 parent (with probability 1 − ℐ_2_). By equating the left and right, we are requiring that, at equilibrium, these class reproductive values do not change from one generation to the next. Solving the two equations above gives

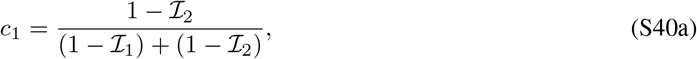

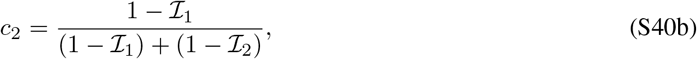

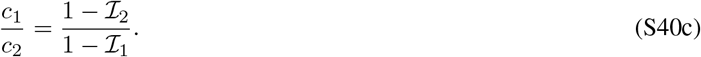

The numbers *c*_1_ and *c*_2_ can equivalently (one would have to solve the same equation) be found using a definition that is common in the literature (see e.g. Taylor, 1990, eq. 8): the vector of class reproductive values [*c*_*f*_ *c*_*m*_]^*T*^ is the left eigenvector associated to the eigenvalue 1 of the matrix ℐ. 1 is always an eigenvalue of this matrix since its row entries sum to 1 along every row.

#### S10.2 Invasion criterion

Now consider that the two classes are in fact the separate sexes, and that most organisms are homozygotes for a resident allele with fitness 1 in comparison to a mutant; we are interesting in the dynamics of this newly arisen rare allele *E*_*s*_. In line with the previous subsection, denote by ℐ_*f*_ the probability that a female-produced gamete i.e. an egg contains her maternally-inherited allele (recall this is all from the perspective of one locus). Similarly, ℐ_*m*_ is the probability that a sperm contains the paternally-inherited allele.

Taking into account the different fitnesses of males and females carrying the rare allele, we can write down a recurrence obeyed by the allele frequency of *E*_*s*_ in eggs and sperms, exactly analogous to Eqs. 21, 19 in the main text. Make only three changes:

- introduce into the equations the fact that an egg contains the maternally inherited allele with probability ℐ_*f*_, and the sperm contains the maternally inherited allele with probability ℐ_*m*_,
- neglect terms that are products of rare-allele frequencies (like in the discussion around Eq. S25) since they will be of negligible value when the allele is rare,
- suppose that any remaining organisms have fitness 1 + *σ*_*m*_ if male and 1 + *σ*_*f*_ if female.

We use the different notation of *σ*_*f*_, *σ*_*m*_ compared to the already-used *s*_*f*_, *s*_*m*_ since we want to suppress the effect of ploidy and dominance in the above remaining organisms (the ones that remain after assuming mutant allele is rare, second bullet point above). For example, when an autosomal allele is rare, it is present mostly in heterozygotes, so *σ*_*i*_ = *hs*_*i*_, the fitness effect of the mutant allele must be weighted by the dominance. However, for an X-linked allele, even when rare, it is haploid in males but diploid in females, so *σ*_*f*_ = *hs*_*f*_, *σ*_*m*_ = *s*_*m*_.

With this in mind, some algebra leads one to recurrences of the same form as Eqs. S30, S31 but including the ℐ terms:

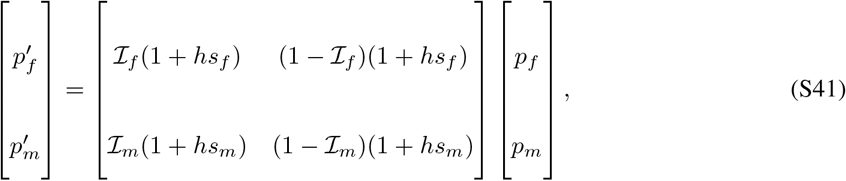

Compare to the matrix equations Eqs. S30, S31. Identical to the derivation of the invasion criteria in the previous section (e.g. S33), the rare allele *E*_*s*_ can invade when the above matrix has dominant eigenvalue larger than one. Applying the formula (S29) to find the dominant eigenvalue and asking when it is larger than 1 exactly as before leads to the invasion criterion

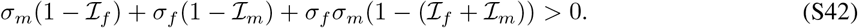

Compare this with Ineq. S33, which has a similar form, and is indeed identical if one notices that for an X-linked locus, ℐ_*f*_ = 1*/*2, ℐ_*m*_ = 0, and *σ*_*f*_ = *hs*_*f*_, *σ*_*m*_ = *s*_*m*_ as already mentioned.

#### S10.3 Combining the two

Looking simultaneously at the invasion criterion Ineq. S42 and the class reproductive values of Eq. S40 shows that they are related. Dividing (S42) throughout by the never-negative quantity (1 − ℐ_*f*_) + (1 − ℐ_*m*_) (that is also found in (S40)) gives us the rephrased inequality

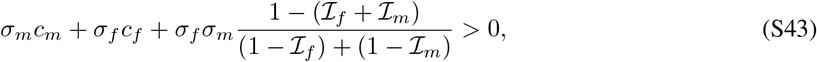

which shows how, and indeed that, class reproductive values affect the invasion criteria.

## Footnotes

1 In the weak selection limit *s*_f_ *s*_m_ ≈ 0 and assuming full dominance *h* = 1, this condition reduces to the reproductive-value weighted criterion *s*_m_ + 2*s*_f_ *>* 0.

